# Proteogenomic analysis reveals adaptive strategies to alleviate the consequences of aneuploidy in cancer

**DOI:** 10.1101/2024.03.05.583460

**Authors:** Jan-Eric Boekenkamp, Kristina Keuper, Stefan Redel, Karen Barthel, Leah Johnson, Angela Wieland, Markus Räschle, Zuzana Storchova

## Abstract

Aneuploidy is prevalent in cancer and associates with fitness advantage and poor patient prognosis. Yet, experimentally induced aneuploidy initially leads to adverse effects and impaired proliferation, suggesting that cancer cells must adapt to aneuploidy. We performed *in vitro* evolution of cells with extra chromosomes and obtained cell lines with improved proliferation and gene expression changes congruent with changes in aneuploid cancers. Integrated analysis of cancer multi-omics data and model cells revealed increased expression of DNA replicative and repair factors, reduced genomic instability, and reduced lysosomal degradation. We identified E2F4 and FOXM1 as transcription factors required for adaptation to aneuploidy *in vitro* and in cancers and validated this finding. The adaptation to aneuploidy also coincided with specific copy number aberrations that correlate with poor patient prognosis. Chromosomal engineering mimicking these aberrations improved aneuploid cell proliferation, while loss of previously present extra chromosome impaired it. The identified common adaptation strategies suggest replication stress, genomic instability, and lysosomal stress as common liabilities of aneuploid cancers.

## Introduction

Aneuploidy, an imbalanced chromosome number that deviates from the multiples of a chromosome set, is a hallmark of cancer found in 90% of solid tumors and 70% of hematopoietic cancers (*1*). This establishes aneuploidy as one of the most common types of genetic alterations in cancer. Yet, experiments conducted in budding yeast, plants, drosophila, and mammalian cell cultures demonstrate that gain or loss of a single chromosome has often detrimental consequences (*2–5*). In an acute response to aneuploidy immediately after chromosome missegregation in human cells, the cells show differential regulation of autophagy and lysosomal stress, metabolic changes, and genomic instability (*6–9*). Human cells with a constitutive gain of an extra chromosome usually show reduced proliferation, conserved changes in gene expression patterns, altered protein homeostasis and genomic instability (*3, 10, 11*). The stresses arise due to a low overexpression of several hundreds of proteins encoded on the extra chromosome, which overloads the protein folding machinery, leading to the accumulation of cytosolic protein aggregates, deregulation of autophagy and increased proteasomal activity (*8, 12, 13*). The genotoxic stress is characterized by reduced expression of key replicative factors, replication stress, the accumulation of DNA damage, and increased occurrence of de novo chromosomal rearrangements (*6, 14, 15*). These characteristic features of aneuploidy document that a gain of an extra chromosome puts a burden on the cellular machineries.

Yet, chromosome copy number changes are frequent in cancer. While some aneuploidies might be random, there are recurrent patterns of copy number changes in cancer genomes suggesting that certain aneuploidies provide the cells with a selection advantage shaping the cancer genome by selection forces (*1, 16–19*). For example, the gains of chromosome arms 8q and 20q are prevalent across many different cancer types (*20*), while loss of chromosome arm 3p is found in squamous cancer, loss of chromosome 10 in glioblastoma, and 13q gain in gastrointestinal tumors (*21*), documenting that some aneuploidies provide specific advantages to cancer cells. Cancer specific aneuploidy patterns are selected likely due to the presence of individual genes, which bring a gene-specific and tissue-specific advantage. Indeed, the number of tumor suppressors and oncogenes on individual chromosomes correlates with the frequency of gains and losses of this chromosome in cancer (*19, 22*). Recent results demonstrate that expression changes due to gain of an individual chromosome might be sufficient to provide a proliferation advantage. For example, gain of chromosome arm 1q in some p53-proficient cancers improves their proliferation, possibly due to the increased expression of MDM4, which suppresses p53 signaling (*23*). Thus, specific cancer aneuploidies, similarly as oncogenic gene mutations, are selected according to their positive effect on cellular fitness (*19, 22*).

Despite the potential advantages associated with the gains of certain chromosomes, the concomitant burden arising from increased expression of hundreds of genes with no direct advantage remains. This is evidenced in engineered aneuploid human cell lines where the impairment of proliferation scales with the number of genes located on the extra chromosome (*24, 25*). Diploid cancer cells tend to maintain more often chromosome arm aberrations, while cancer cells that underwent whole genome doubling can tolerate whole chromosome aberrations (*26*). In cells with elevated chromosomal instability (CIN+) induced by spindle assembly checkpoint inhibitors, survivors with complex karyotypes arise, but these cells become soon outcompeted by cells carrying smaller, more precise copy number alterations (*16*). Thus, even if a chromosome gain provides a selective advantage to the cells, they must concomitantly adapt to adverse effects of excessive genetic material.

The mechanisms by which cells adapt to tolerate or counteract the deleterious consequences of chromosome gain remain elusive. Here, we investigated the mechanisms that enable human cells and cancers to adapt to aberrant karyotype. To this end, we conducted *in vitro* evolution experiments, where constitutive polysomic cell lines were passaged over an extended period. The polysomic cell lines displayed improved proliferation after *in vitro* evolution while maintaining the extra chromosome. The improved proliferation of aneuploid cells was associated, among others, with reduced DNA damage and genomic instability, reduced replication stress, and an altered expression of factors involved in DNA replication and lysosomal degradation. The identified pathways and genes tightly correlate with pathways deregulated in aneuploid cancers. The gene expression changes in model cells aligned with gene expression changes observed in aneuploid cancers and uncovered the transcriptional factors E2F4 and FOXM1 to critically contribute to adaptation to chromosome gains, which we experimentally validated. Moreover, we demonstrate that specific chromosome 5 aberrations that alleviate the consequences of aneuploidy *in vitro* also correlate with poor patient prognosis. Our integrated approach elucidates the molecular landscape of adaptation to aneuploidy and identifies potential vulnerabilities in aneuploid cancers.

## Results

### Cells with additional chromosomes improve their proliferation over time

To study adaptive changes occurring during the evolution of cells harboring defined chromosome gains, we used newly prepared aneuploid cell lines into which chromosome 5 or chromosome 21 was introduced by microcell-mediated chromosome transfer (MMCT), as previously described (*3, 25*). Two cell types, HCT116 and RPE1, were used to generate polysomic cells with a trisomy of chromosome 5 (hereafter Htr5 and Rtr5), a tetrasomy of chromosome 5 (Hte5), or a trisomy of chromosome 21 (Rtr21). Gain of an extra chromosome leads to cellular stress manifested by reduced proliferation compared to the parental control (*3*). To elucidate how human cells adapt to chromosome gain, we cultured the cell lines in a nutrient-rich medium for 50 passages in biological triplicates (p50, approximately 150 generations, Fig. 1A). The medium lacked the antibiotics required for the selection of the extra chromosome, allowing for the potential loss of any chromosome. After the *in vitro* evolution, almost all polysomic cells exhibited a significant increase in proliferation, quantified by the increased area under the growth curves (AUC), in comparison to the original, p0 population (Fig. 1B, Supplementary Fig. 1A-C, Supplementary Table 1). The increase was observed in 10 out of 12 independent replicates, regardless of the specific cell line or the added chromosome, albeit to a varying degree (Supplementary Fig. 1C). In contrast, the proliferation rate of the parental diploid cell line remained nearly unchanged (Fig. 1B, Supplementary Fig. 1A, C). Despite the significant increase in proliferation, all evolved polysomic clones exhibited less efficient proliferation than the parental disomic cells (Fig. 1B, Supplementary Fig. 1A). Thus, human cell lines adapt to chromosome gains after culturing over extended periods of time to counteract the adverse effects that initially caused their poor proliferation.

**Figure 1.**
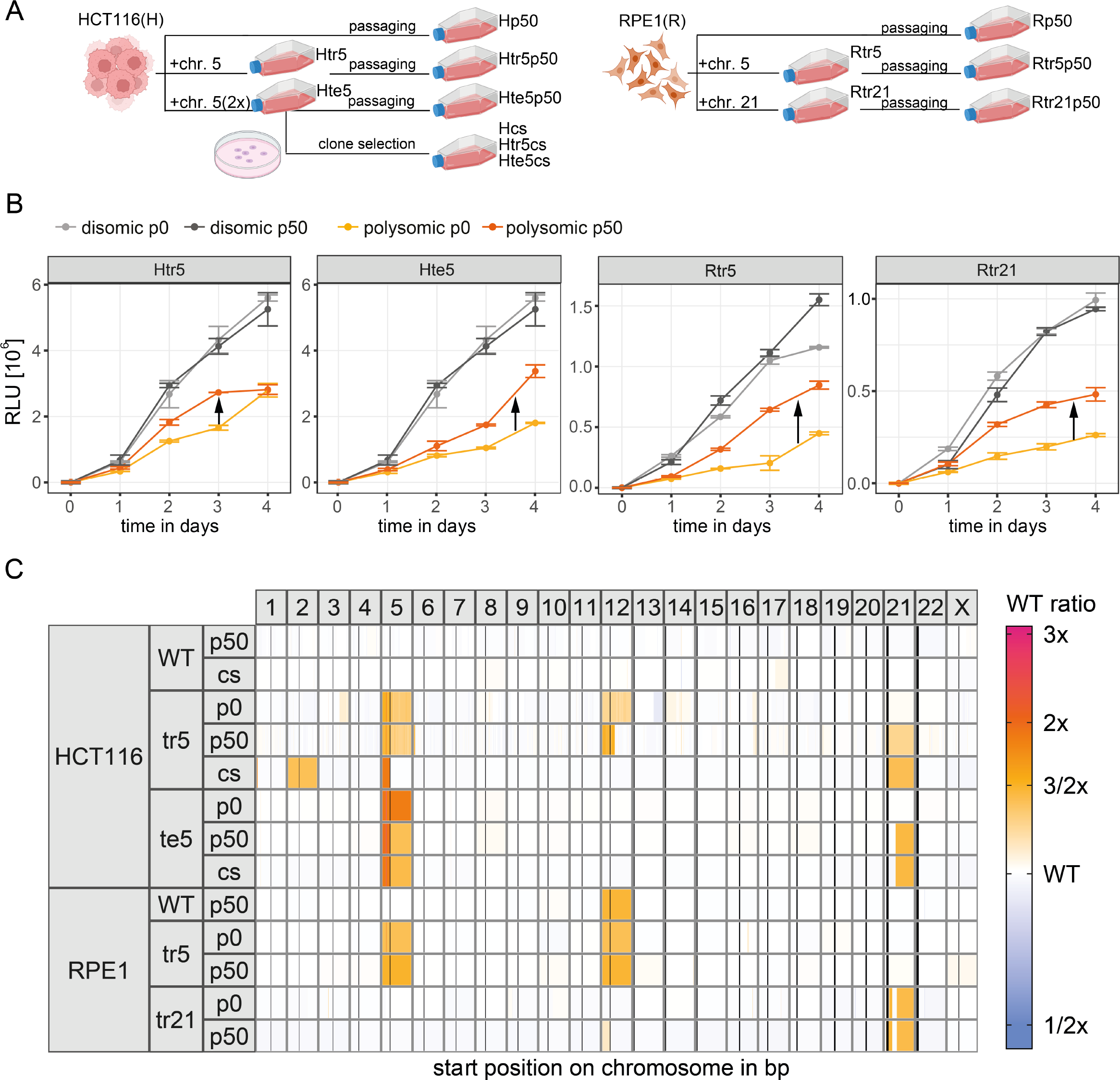
Improved proliferation following *in vitro* evolution of polysomic human cells. **A**. Schematic illustrating the experimental setup of *in vitro* evolution applied to polysomic model cell lines. **B**. Population growth of the cell lines before and after *in vitro* evolution evaluated by MTT assay and normalized to the time point 0. Points represent mean relative light units (RLU) with SEM. **C**. Chromosome copy number ratios relative to parental, unevolved WT for each cell line (row) and chromosome (column). Vertical bars within chromosomes depict the location of centromeres.

### Loss of the additional chromosome is not a pre-requisite for improved proliferation in polysomic cells

To further characterize the adaptive processes that help aneuploid cells to overcome the initial growth inhibition we subjected the evolved clones to an in-depth karyotype analysis. From each cell line, we selected the clonal lines that showed the highest increase in proliferation compared to the unevolved p0 cell line (Fig. 1B and S1A) and subjected them to either low-coverage whole genome sequencing (WGS) or array-based comparative genomic hybridization (aCGH) (Supplementary Table 2). The *in vitro* evolved clones maintained the extra chromosomes, and additional copy number changes affecting chromosome arms, as well as smaller regions, were observed (Fig. 1C, Supplementary Table 2). For instance, we identified sub-clonal losses of the q arm of chromosome 5 in both Htr5 and Hte5 cell (Fig. 1C). Additionally, full or partial gains of chromosome 21, and less frequently 2, were observed in polysomic cell lines after in-vitro evolution (Fig. 1C). The trisomic cell lines derived from non-cancerous RPE1 cells maintained the extra chromosomes as well and displayed no clonal changes in their karyotypes, in line with the previous observation that polysomy leads to only mild genomic instability in RPE1 cells (*14*). The parental RPE1 gained an additional chromosome 12 and showed some characteristics of trisomic cells (Fig. 1C, see below). The gain of chromosome 12 is in line with previous observation that RPE1 cells tend to gain chromosome 12 (*27*). Based on these data we conclude that the improved proliferation did not occur simply due to the loss of the extra DNA. Indeed, we found that the total amount of altered DNA in the adapted polysomic cells relative to WT only weakly correlates with the observed proliferation improvement (Supplementary Fig. 1D).

In an alternative approach, we plated the HCT116-derived cells at a low density and selectively collected the largest, best proliferating colonies originating from a single cell (referred to as colony selection or “cs” Fig. 1A). These cells showed similar chromosomal copy number changes (Fig. 1C) suggesting that the cells with these specific aberrations were present in the original population and the prolonged passaging allowed their outgrow.

Cell proliferation can be enhanced through reduced duration of the phases of the cell cycle, or decreased rates of senescence or cell death. Cells after chromosome gain exhibit prolonged G1 phase duration and subtle changes in the percentage of senescent and dead cells compared to the parental cell lines. We found that the fraction of cells in the G1 phase decreased, while fraction of cells in the S phase increased following in vitro evolution in polysomic clones (Supplementary Fig. 2A, B). There were no uniform changes in the fraction of dead and senescent cells within the population which could fully explain the proliferation improvement (Supplementary Fig. 2C, D). Moreover, the p53 abundance and signaling was not changed in the evolved clones (Supplementary Fig. 2E). We conclude that the in vitro evolved polysomic cell lines improved their proliferation despite maintaining the extra chromosome and functional p53 signaling. Thus, the obtained clones provide a valuable model for identifying the physiological changes required for adaptation to chromosome gain.

### Analysis of gene expression changes reveals differentially regulated pathways in evolved polysomic cells

To elucidate the molecular mechanisms that enable the enhanced proliferation of the adapted polysomic cells, we analyzed the global proteome before and after *in vitro* evolution by quantitative mass spectrometry using a tandem mass tag (TMT)-based quantification strategy (Supplementary Table 3) (*28, 29*).These results were then compared to aneuploidy-specific expression signatures extracted from the TCGA and CPTAC pan-Cancer data repositories by independent bioinformatic analysis (Fig. 2A, see below).

**Figure 2.**
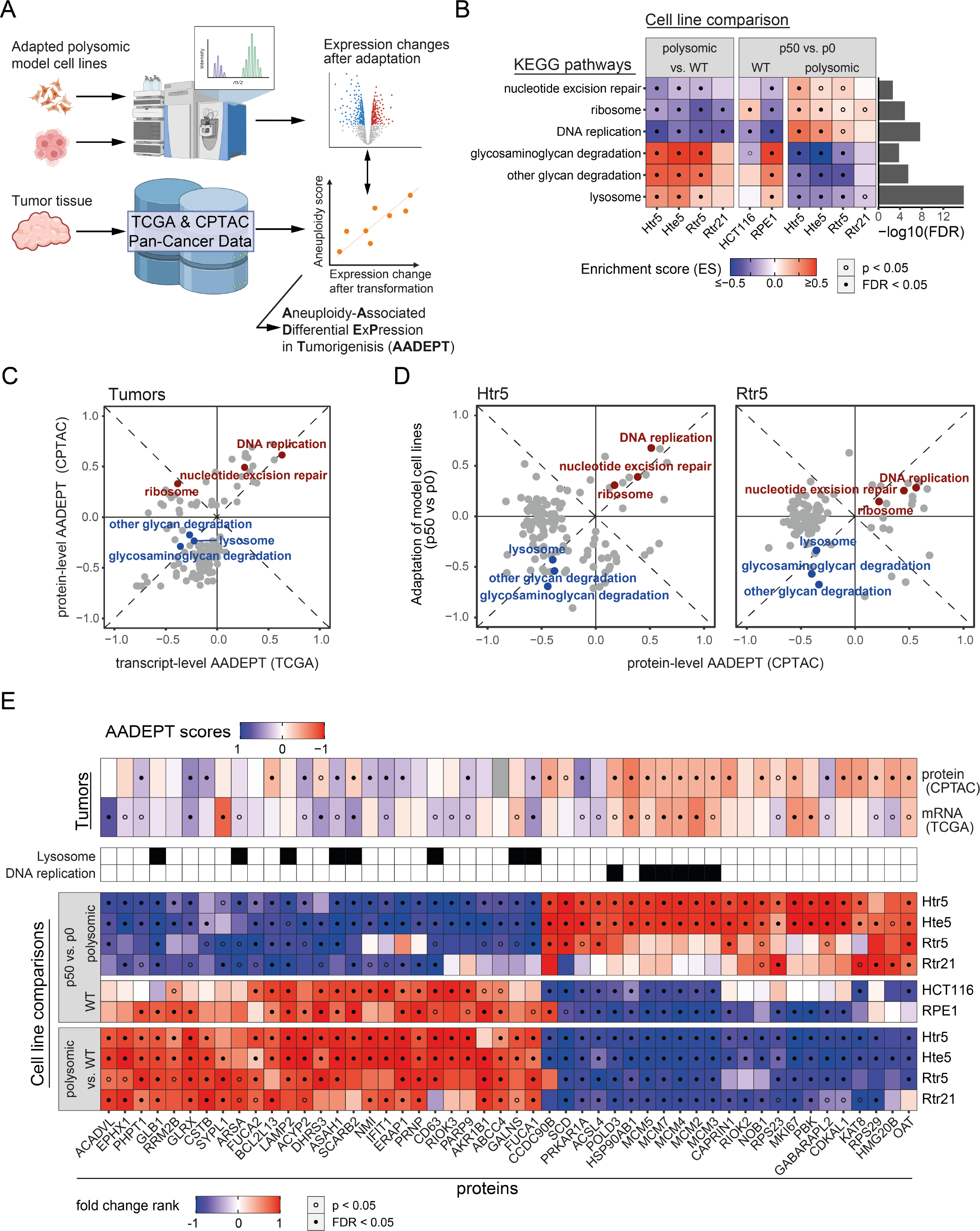
Expression changes in genes and pathways after *in vitro* evolution of polysomic human cell correlate with changes in human aneuploid cancers. **A**. Schematic depiction of the data analysis pipeline including the calculation of correlations between aneuploidy scores and gene expression changes after transformation (primary tumor vs. normal tissue) in TCGA and CPTAC tumor samples – AADEPT (Aneuploidy-Associated Differential ExPression in Tumorigenesis). **B**. Enrichment scores of pathways significantly altered in at least three polysomic cell lines after *in vitro* evolution. The statistical significance of fold change enrichment in individual comparisons is indicated by the points within each tile. FDR bars (cut-off at 0.05) are based on multivariate ANOVA for overall enrichment between p50 vs. p0 comparisons. **C**. 2D pathway enrichment of AADEPT scores on transcript-(TCGA) and protein-level (CPTAC), showing enrichment scores for pathways with FDR < 0.05 and highlighting those with shared enrichment in adapted polysomic cell lines (blue - negative, red - positive). **D**. 2D pathway enrichment comparing protein-level AADEPT scores with fold changes in polysomic Htr5 and Rtr5 after adaptation, showing enrichment scores for pathways with FDR < 0.05. **E**. Top 50 proteins according to their relevance score, which shows the correlation of their AADEPT scores (above) and their relative abundance changes in adapted polysomic model cell lines (below). Fold changes are ranked and scaled from -1 (lowest) to 1 (highest).

Normalizing the protein abundance values to the disomic parental cell lines to obtain log2 fold changes (FC) and visualizing them grouped by chromosome identity (Supplementary Fig. 3A, Supplementary Table 3) confirmed the increased expression from the extra chromosomes in evolved cell lines. Abundance of proteins encoded on the supernumerary chromosome should be increased by 50 % for trisomic cells and doubled in cells with two additional chromosome copies. However, previous work revealed gene dosage compensation, which reduces the abundance of some proteins towards levels measured in diploids (*3, 30*). Efficient dosage compensation might reduce the burden on polysomic cells and thus improve their proliferation without loss of the additional chromosome. Comparing the degree of dosage compensation in our polysomic cells, we found an increase for six chromosome arms in *in vitro* evolution experiments, while there was a reduction or no change in two cases, leading to a slight, but non-significant, increase of dosage compensation after *in vitro* evolution (Supplementary Fig. 3B, C). We conclude that the improved proliferation in adapted polysomic cells cannot be explained entirely by enhanced dosage compensation of genes located on the supernumerary chromosomes.

We next applied standard differential expression analysis to identify genes whose expression changes show similarity among polysomic cell lines. Upon chromosome gain, at p0, we identified 55 commonly downregulated and 40 upregulated proteins (FDR < 0.05, Supplementary Fig. 4A), confirming previous observations that chromosome gain elicits gene expression dysregulation independently of the gained chromosome or the cell line (*3, 31*). In contrast, the paths to adaptation to polysomy are less convergent, as only two upregulated and three downregulated factors were shared among all evolved polysomies, but not in the parental cell lines (Supplementary Fig. 4A). These were GLB1 (encoding lysosomal hydrolase galactosidase beta 1), CTSA (encoding lysosomal peptidase cathepsin Z) and BCL2L13 (inhibitor of apoptosis from the BCL2 family), whose abundance was commonly decreased with adaptation, while PRPF19 (coding for ubiquitin-protein ligase involved in splicing and DNA repair) and OAT (encoding mitochondrial ornithine aminotransferase), were more abundant after evolution (Supplementary Fig. 4B).

With this low turnout of individual proteins, we sought a better insight into the functional changes by using the fold changes of relative protein abundances between p50 and p0 to test metabolic pathways for significant enrichment (KEGG (*32*)). This analysis revealed a set of six pathways including DNA replication, nucleotide excision repair (NER) and ribosome, which became downregulated upon chromosome gain, but showed increased expression following *in vitro* evolution in both, HCT116 and RPE1 derived polysomic cell lines (Fig. 2B, Supplementary Table 4, see Material and Methods). Additionally, lysosomal pathways essential for the degradation and recycling of biomolecules, were upregulated after the addition of a chromosome, but showed a marked downregulation after 50 passages (Fig. 2B). Thus, we identified pathways changes associated with cellular response to aneuploidy that were reversed after *in vitro* evolution independently of the parental cell line and added chromosomes.

### Adaptations to aneuploidy *in vitro* correspond with the adaptations in aneuploid cancer

We next aimed to leverage our model of adapted polysomic cell lines together with publicly available multiomics data of patient-derived aneuploid tumors to identify molecular mechanisms that facilitate the adaptations to aneuploidy in an integrative bioinformatic analysis. To this end, we used the transcriptome data from the TCGA PanCanAtlas (*28, 33*) and the proteome data from the CPTAC (*29*) for both cancerous and adjacent non-cancerous tissues, as well as the corresponding genomic copy number data. We then calculated the difference in gene expression between primary tumors and the respective patient’s normal tissue and correlated the expression changes with the tumor’s aneuploidy score (AS), which was derived as a sum of altered chromosome arms (*21*) (Supplementary Table 4). With this approach, we obtained a comparative measure called **AADEPT** (**A**neuploidy-**A**ssociated **D**ifferential **E**x**P**ression during **T**umorigenesis), which quantifies the cellular adaptations in cancer transcriptome and proteome of malignant tumors linked to varying degrees of aneuploidy (Supplementary Fig. 5A, B, Supplementary Table 5). Comparing the calculated AADEPT scores from transcript-(TCGA) and protein-level (CPTAC) showed a striking correlation of pathway enrichments, with DNA replication and nucleotide excision repair being upregulated and lysosomal pathways being downregulated (Fig. 2C). Most strikingly, all pathways that were universally deregulated in the evolved polysomic model cells correlated with the AADEPT in transformed aneuploid tissues except for ribosomes on transcript-level. The quantified AADEPT scores also correlated strongly with the changes in evolved polysomic cell lines, including an overlap in overabundance of RNA polymerase, spliceosome, and DNA repair proteins, and reduced abundance in lysosome and lysosomal degradation pathways (Fig. 2D, Supplementary Fig. 6A). Finally, we explored a possible overlap with the hallmark gene sets (MSigDB (*34*)) and its enrichment with AADEPT scores. Again, we observed a striking level of congruence with the protein abundance changes in adapted model cells, with a stronger similarity observed in the polysomic cell lines derived from cancerous HCT116 (Supplementary Fig. 6B). Specifically, E2F target proteins, MYC targets and G2/M checkpoint proteins were more abundant, whereas innate immune response proteins reduced upon adaptation to aneuploidy in both model cell lines and malignant tumors. Thus, the significantly differentially regulated pathways after *in vitro* evolution of polysomic cells strongly correlate with the differentially regulated pathways in aneuploid tumors.

The pathway analysis revealed that the common expression changes in evolved polysomic cell lines overlap with pathway changes in aneuploid cancers. Therefore, we reevaluated the protein-level changes shared among evolved polysomic cell lines, this time using a less conservative approach. To this end, we merged the proteome fold change values to calculate for each protein a “relevance score”, which represents the expression changes between p50 vs. p0, and between polysomic vs. parental cell lines at p0. Moreover, this score penalizes the changes in protein abundance occurring during the evolution of wild type cells to emphasize the aneuploidy-specific adaptations (Supplementary Fig. 5C, Supplementary Table 3, Material and Methods). By this approach, we determined the top 50 proteins representative of the reversion of aneuploidy-induced expression changes after *in vitro* evolution observed on pathway-level. Strikingly, many of these changes correspond with the AADEPT score on both transcript- and protein-level in TCGA and CPTAC patient samples (Fig. 2E). For example, we identified upregulation of several replication proteins (five out of the six subunits of the MCM2-7, or DNA polymerase delta 3), upregulation of heat shock protein HSP90, and downregulation of factors related to lysosome function (LAMP2, CSTB) as well as factors related to interferon response (e.g. IFIT1). Of note, the degree of changes in the non-cancerous RPE1 models is generally lower in both the multivariate pathway enrichment (Fig. 2B) as well as in the protein relevance analysis (Fig. 2E). Based on these data, there are common pathways and proteins that emerge as decisive factors influencing proliferation in aneuploid cells. Consequently, these pathways may constitute the central anti-proliferative mechanisms associated with chromosome gains. Taken together, our computations meta-analysis identified the key pathways altered during adaptation to aneuploidy *in vitro* and during tumorigenesis.

### Restoration of DNA replication and repair processes reduce genomic instability in polysomic clones after *in vitro* evolution

Among the most prominent changes observed upon *in vitro* evolution was the restored abundance of proteins that are assigned to “DNA Replication” (KEGG), which are generally downregulated upon chromosome gain (Fig. 3A) (*7, 14*). Many of the identified proteins were also identified by their high AADEPT scores in TCGA and CPTAC (Fig. 3A). Among the most affected proteins were the subunits of the MCM2-7, a heterohexameric complex required for replication licensing, which together with CDC45 and GINS complex acts as a replicative helicase during DNA replication (so-called CMG), which we confirmed by immunoblotting of the MCM2-7 subunits in non-evolved and evolved polysomic cells (Fig. 3B, C, Supplementary Fig. 7A). No changes in MCM2-7 expression were observed in the HCT116 (note that RPE1p50 gained chromosome 12 and was not used for further analysis, Fig. 1B). We asked whether the changes in abundance of replication factors have an impact on the dynamics of DNA replication. To this end, we used DNA combing and visualized full track lengths of labeled replicating DNA (Supplementary Fig. 7B). Quantification of the replication rate revealed a reduction upon *in vitro* evolution in Rtr21 and Hte5 (Supplementary Fig. 7C). Next, we measured the inter-origin distance, where we observed a significant increase in the inter-origin distance of Hte5 cell line, carrying the highest amount of additional DNA, which was then reduced upon evolution (Supplementary Fig. 7D). Since replication stress often results in disturbances of the fork symmetry, we measured the neighboring fork lengths. We found that the percentage of asymmetric forks is increased in cells with an additional chromosome, but this percentage decreased after evolution (Fig. 3D), suggesting reduced replication defects after *in vitro* evolution.

**Figure 3.**
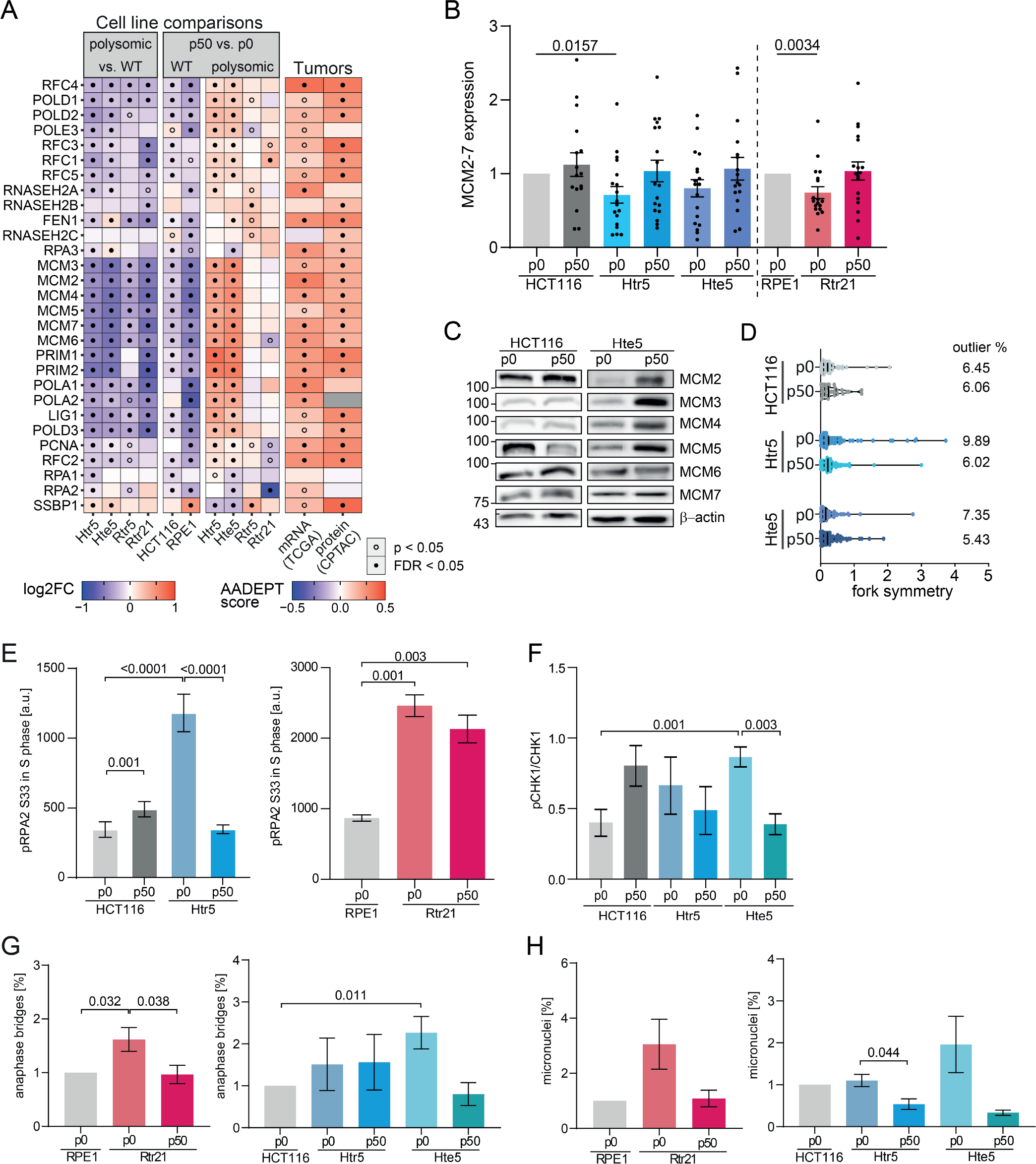
Increased expression of replicative factors and reduced replication stress, genomic instability, and DNA damage after *in vitro* evolution. **A**. Expression changes of factors involved in replication after chromosome gain (polysomic vs. WT) and after *in vitro* evolution (p50 vs. p0) and their comparison with the AADEPT scores (as in Fig. 2E). **B**,**C**. Immunoblotting of the MCM2-7 subunits and the quantification in the model cell lines. The values for all subunits were normalized to the respective parental cell line and pooled together (N: 16 – 18, each MCM subunit at least 2x). **D**. Fork symmetry in the model cell lines before and after evolution and the 0.5 percentile of outliers (identified with ROUT). At least three independent experiments were performed, N ≥ 33 in each experiment. **E**. Median of relative phosphorylation of the RPA2 on the S33 during S-phase evaluated by flow cytometry. Three biological experiments performed in 1-2 technical replicates are shown. **F**. Quantification of relative phosphorylation of CHK1 on S345 evaluated by immunoblotting. Three biological experiments. **G**. Quantification of the occurrence of anaphase bridges in model cell lines. At least three independent experiments (N: 3 - 12), at least 30 cells in each were evaluated. **H**. Quantification of the occurrence of anaphase bridges in model cell lines. At least three independent experiments (N: 3 - 12), at least 50 anaphase cells in each experiment were evaluated. Scale bars in microscopy images 5 Um. Data information: Mean with SEM is shown in all experiments. P-values were calculated using unpaired Student’s t-test.

Evaluation of the replication protein RPA2, which becomes phosphorylated at RPA2 S33 by the ATR kinase upon replication stress, showed that the phosphorylation in the S-phase increases upon chromosome gain, in line with our previous findings (*14*). Importantly, the phosphorylation of RPA2 was reduced in evolved polysomic cell lines (Fig. 3E). Similar reduction was observed in the phosphorylation of CHK1 S345, another substrate of the ATR kinase (Fig. 3F). Next, we asked whether the altered gene expression and partly reduced replication stress observed in evolved cells affect genomic instability. Occurrence of anaphase bridges, which result from defective DNA repair or replication, is increased upon chromosome gain, but was reduced in evolved Hte5p50 and Rtr21p50 cell lines compared to the respective p0 (Fig. 3G). Similarly, accumulation of micronuclei, another marker of genomic instability, was reduced upon *in vitro* evolution (Fig. 3H). We conclude that one of the strongest changes upon *in vitro* evolution of cells with additional chromosomes is an increased expression of DNA repair and replication factors and reduced genomic instability, which is reflected by reduced checkpoint signaling and decreased DNA damage.

Our analysis also revealed a reduction of lysosomal gene expression in the evolved aneuploid cells, such as lysosomal cathepsins CTSA and CTSB, LAMP1 and 2 (Lysosomal-associated membrane protein 1 and 2) and others (Supplementary Fig. 8A). It should be noted, however, that the AADEPT scores of these proteins were more heterogenous, suggesting a weaker overlap with gene expression changes in aneuploid cancers. Immunofluorescence imaging of LAMP1 revealed fewer foci in the evolved cells, while lysotracker and LAMP2 analysis indicates rather mild and heterogenous effects (Supplementary Fig. 8B, C). We conclude that while abundance of lysosomes and lysosomal proteins likely decreases upon evolution of aneuploid cells, but further studies will be required to characterize the observed changes.

### FOXM1 and E2F4 transcription factors contribute to improved proliferation in evolved polysomic cells

To get more insight into the molecular pathways responsible for the improved proliferation, we asked which transcription factors orchestrate the gene expression changes observed in polysomic cells after *in vitro* evolution. We focused on HCT116 derived polysomic cell lines, as there was strong significant correlation between the evolution of polysomic cancer cell lines derived from HCT116 and AADEPT scores calculated from the TCGA and CPTAC data (Fig. 2D, E). We determined the most differentially regulated proteins after *in vitro* evolution in Htr5 and Hte5 (FC > 3/2 or FC < 2/3, FDR < 0.05) and specifically excluded proteins with evidence for corresponding, significant abundance changes in the evolution of the WT cell lines. The 43 identified proteins exhibited the pattern characterized by the reversion of expression changes with the adaptation of our polysomic HCT116 cells (Fig. 4A). Ten of these proteins were upregulated after chromosome gain and downregulated following *in vitro* evolution. Among them, we again found factors involved in interferon response and integrin signaling, such as IRF9 (transcription factor that mediates innate immune response), ITGA3 (integrin subunit alpha 3 is a receptor involved in cell migration) or RBCK1 (E3 ubiquitin ligase, which regulates the inflammatory response) and other factors involved in signaling by type I interferons, strongly overlapping with the global meta-analysis (Fig. 4A). This suggests, together with the comparison with hallmark gene signature enrichment (Supplementary Fig. 6B), that downregulation of the inflammatory response contributes to improved proliferation of aneuploid cells.

**Figure 4.**
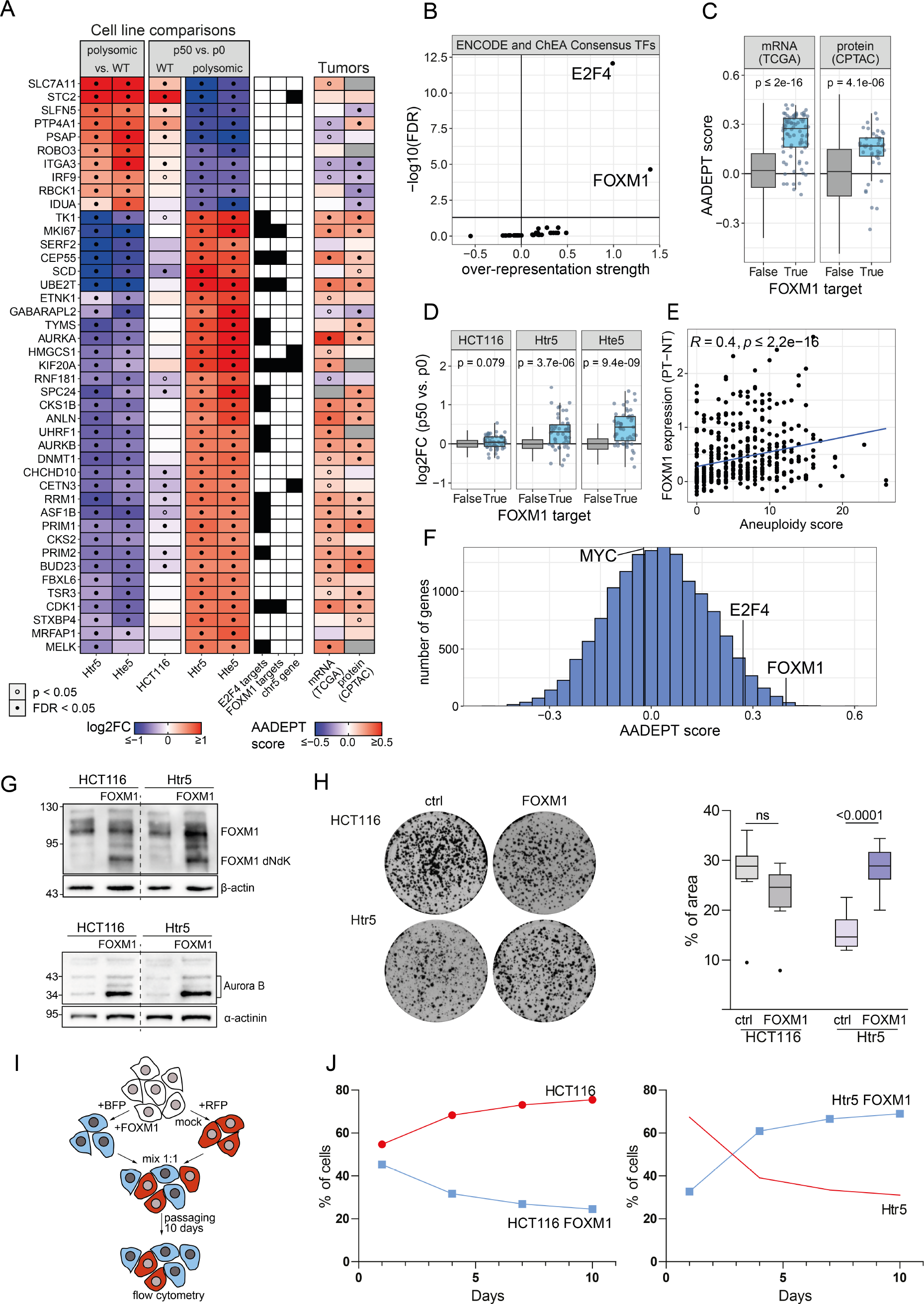
FOXM1 overexpression rescues proliferation defects in cells with extra chromosomes. **A**. Top 42 proteins with fold changes higher than 3/2 or lower than 2/3 before and after *in vitro* evolution of HCT116-derived polysomic model cell lines, and the corresponding AADEPT scores. **B**. Over-representation of ENCODE and ChEA Consensus transcription factor (TF) targets among the differentially regulated genes in the evolved polysomic cell lines and aneuploid tumors. **C**. Transcript-(TCGA) and protein-level (CPTAC) AADEPT scores of FOXM1 target genes were tested against all other proteins using Welch’s t-tests (N: 82, N: 42). **D**. Protein abundance fold changes of all FOXM1 targets in evolved model cell lines, tested against all other proteins using Welch’s t-tests (N: 56). **E**. Spearman correlation coefficient between TCGA patient tumor aneuploidy score and FOXM1 gene expression relative to normal tissue (N: 467). **F**. Histogram of transcript-level AADEPT scores derived from TCGA with highlighted scores of MYC, E2F4, and FOXM1. **G**. Representative immunoblot of overexpression of FOXM1 and FOXM1 targets in the parental HCT116 and unevolved Htr5. **H**. Representative images of clonogenic assay of HCT116 and Htr5 with and without overexpression of FOXM1, with quantification of the percentage of area covered by cells (N: 9), mean with quartiles is shown, unpaired Student’s t-test. **I**. Schematic depiction of the competition assay. **J**. Quantification of the RFP and BFP positive cell fraction in competition assay during the incubation period.

Among the top proteins that were downregulated after chromosome gain, but upregulated upon adaptation, we found many cell cycle proteins, such as CDK1, CEP55, MKI67, CDC20, KIF20A, PRIM1, and UBE2T (Fig. 4A). These proteins are all essential for various aspects of cell cycle regulation and cell division, and are frequently dysregulated in cancer (*35*). Next, we identified an overlap between the top upregulated proteins after *in vitro* evolution and the factors with high AADEPT scores in the TCGA and CPTAC to determine the significant over-representation of consensus transcription factor target sets from ENCODE and ChEA. This analysis also revealed a significant enrichment of E2F4 and FOXM1 targets in evolved cancer cell lines and in aneuploid tumors (Fig. 4B-D). Additionally, the expression of FOXM1 itself significantly positively correlates with aneuploidy in cancer (Fig. 4E). In fact, FOXM1 is among factors with the highest AADEPT score (Fig. 4F). Expression of the E2F4 targets and E2F4 also show high AADEPT score, although to a lesser degree than FOXM1 (Fig. 4F, Supplementary Fig. 9A-C). In contrast, the expression of the transcription factor MYC, another pro-proliferative TF, does not correlate with aneuploidy. We conclude that the targets of the transcription factors FOXM1 and E2F4 are pivotal in mediating the adaptation to aneuploidy.

We were particularly interested in the transcription factor FOXM1, which has been associated with aneuploidy previously (*36–38*). Immunoblotting revealed that the FOXM1 expression did not increase, suggesting that rather the regulation of FOXM1 was altered after *in vitro* evolution (Supplementary Fig. 9D). Indeed, the initially decreased levels of Aurora B kinase, a target of FOXM1, were increased in the trisomic cells after *in vitro* evolution (Supplementary Fig. 9E). To directly test the involvement of FOXM1 in proliferation of aneuploid cells, we overexpressed the constitutively active truncated form of FOXM1 (FOXM1-dNdK) in the parental HCT116 and in Htr5 (p0, before evolution). Immunoblotting revealed a strong expression of FOXM1 and increased expression of its targets, such as Aurora B (Fig. 4G, Supplementary Fig. 9F-H). High abundance of FOXM1-dNdK resulted in a significant improvement of proliferation in Htr5 cells, but not in the parental HCT116 (Fig. 4H, I). Next, we performed a competition assay to test whether cells overexpressing FOXM1 will gain an advantage in a mixed population. Here, we mixed HCT116 or Htr5 cells expressing BFP and FOXM1 with cells expressing RFP and a control plasmid, and the fluorescence ratio was measured over 10 days of passaging (Fig. 4I). As expected, the unevolved polysomic cells overexpressing FOXM1 outgrew polysomic control cells, while FOXM1 overexpression rather reduced the proliferation of parental HCT116 (Fig. 4J). Similar results were obtained when the BFP and RFP staining was reverted (Supplementary Fig. 9J). Thus, chromosome gain results in a reduced expression of pro-proliferative FOXM1 targets. Increased expression of these proteins, either after *in vitro*evolution or by FOXM1 overexpression, enables proliferation improvement in trisomic cells, but not in cells with normal ploidy.

### Loss of chromosome arm 5q positively affects proliferation of cell lines with trisomy 5

One of the striking observations following evolution of cells with extra chromosome 5 in HCT116 cell line was the frequent loss of the 5q, while maintaining the 5p arm after *in vitro* evolution (Fig. 1C). Chromosome 5 is a frequent target of large copy number alterations in several malignancies, such as ovarian, gastric, and oesophageal cancer, and malignant myeloid diseases (*18*). Moreover, analysis of chromosome arm level events in the TCGA dataset clearly shows that loss of chromosome arm 5q and gain of 5p are among the most frequent events (Fig. 5A). We asked whether these specific changes in copy numbers of chromosome 5 – loss of 5q with simultaneous retention of 5p – could affect the proliferation of the evolved cells. To this end, we used the recently developed technique ReDACT-TR (Restoring Disomy in Aneuploid cells using CRISPR Targeting with Telomere Replacement (*23*)), and transfected cells of a separate clone of Htr5p0 (before evolution) with a gRNA that cuts near the centromere of chromosome 5, simultaneously with a cassette encoding 100 repeats of the human telomere seed sequence (Fig. 5B). Targeting the q-arm generated two independent cell lines with 5p trisomy, which were confirmed by FISH with probes specific for 5p and 5q arms and by shallow WGS (Fig. 5C, D). No cell lines with trisomy of the q-arm were generated, but we have obtained two diploid cell lines, most likely due to the CRISPR/Cas9 induced loss of chromosome 5 (*39, 40*). Clonogenic assay showed that the trisomy of 5p accompanied by deletion of the 5q-arm resulted in an improved proliferation in both clones Htr5p 1 and Htrp5p 2 compared to the full trisomy of chromosome 5 (Fig. 5E). The ReDACT-TR also generated two clones with restored disomy of chromosome 5, Hdi5 1 and Hdi5 2 (Fig. 5C, D). The elimination of the supernumerary chromosome should theoretically lead to improved cellular proliferation, given that the original defect - chromosome gain - has been rectified. Intriguingly, the proliferation of both Hdi5 clones was significantly impaired, even when compared to the parental trisomy (Fig. 5E).

**Figure 5.**
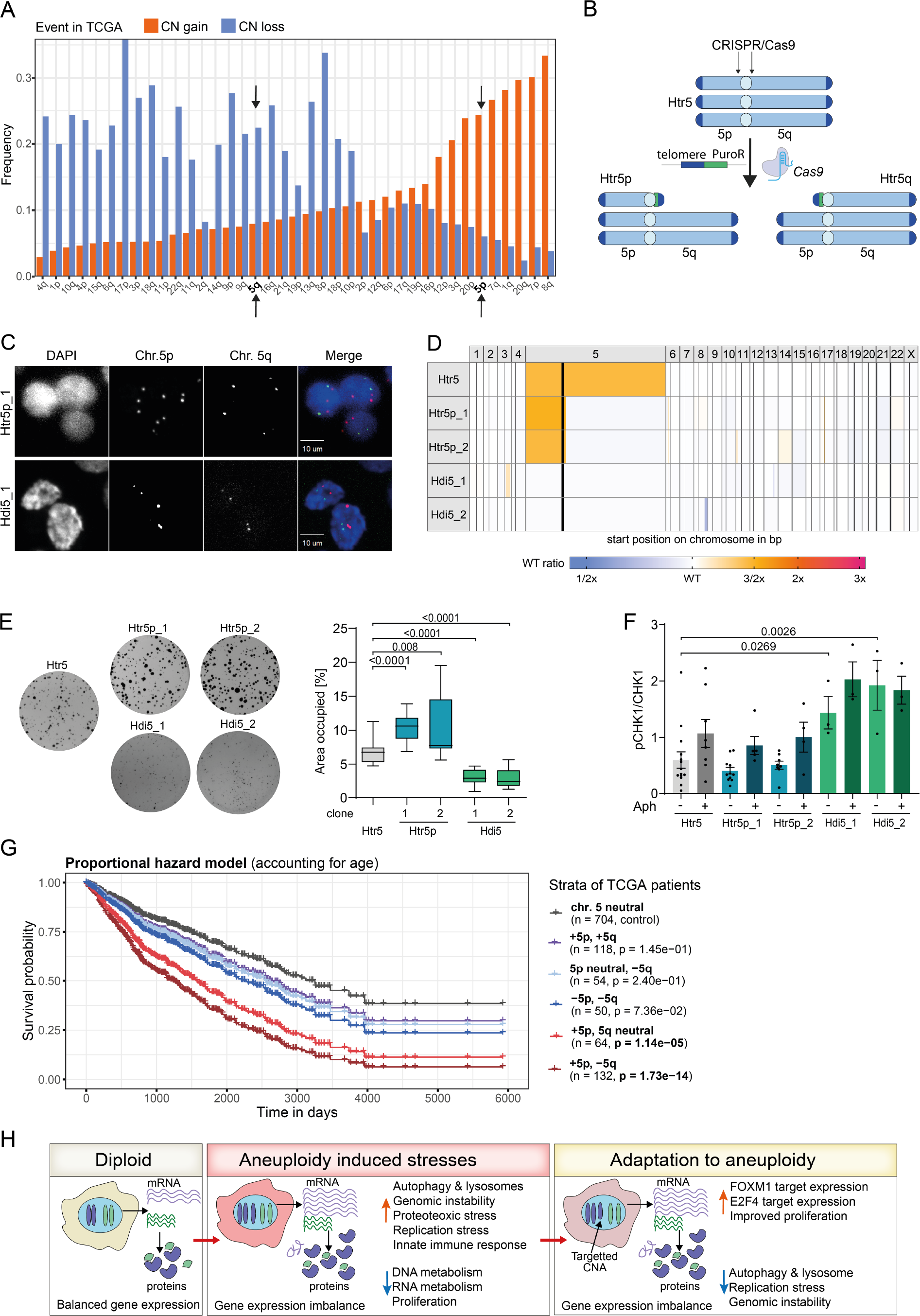
Chromosome 5p arm shows a strong effect on cellular proliferation. Frequency of chromosome arm gains and losses in cancer calculated from TCGA data. Black arrows indicate the frequent loss of 5q and frequent gain of 5p. **B**. Schematic depiction of the chromosome engineering using ReDACT-TR. **C**. Representative images of the fluorescence *in situ* hybridization of probes against 5p (red) and 5q (green) arms. **D**. Chromosome-assigned shallow WGS of the parental Htr5 with Htr5p and Hdi5 cell lines. **E**. Representative images of clonogenic assay with quantification of the area occupied by colonies. For Htr5 and Htr5p cell lines, 16 data points were obtained from three biological replicates at 4 – 6 technical experiments each, for Hdi5 cell lines, 12 data points were obtained from two biological replicates at 4 – 6 technical experiments each. Significance was evaluated using Student’s t-test. **F**. Mean pCHK1(S345)/CHK1 ratio determined via Western Blot (Biological replicates ≥ 3; 1-3 technical replicates each; Student’s t-test), aphidicolin (Aph) treatment was performed for 24 h, 200 nM. **G**. Survival curves of TCGA tumor patients stratified by the cooccurrence of arm-level chromosome 5 copy number gains and losses as fitted by a Cox proportional hazard model, showing the predicted survival curves of a patient of average age (60.7 years). P-values are derived from Wald tests and represent the statistical significance of the hazard ratios relative to patients with copy number neutral chromosome 5. **H**. Model of adaptation to aneuploidy in human cell lines and tumors.

Next, we asked what the underlying molecular mechanisms associated with the proliferative advantages of chromosome 5 q-arm deletion are. Since our previous data showed largely changes in replication factors and DNA damage to affect proliferation of the aneuploid cells, we analyzed the expression of replicative factors, DNA damage and replication stress signaling, which were most differentially regulated after *in vitro* evolution. We observed no changes in MCM5 and MCM7 abundance in the Htr5p clones, as well as no reduction in DNA damage, as evaluated by yH2AX signal and RPA2 S33 phosphorylation (Supplementary Fig. 10A-C). Phosphorylation of CHK2, an ATM-dependent marker for DNA-damage, was also not altered (Supplementary Fig. 10C). In contrast, the cell lines lacking an extra copy of chromosome 5q arm showed slightly reduced CHK1 phosphorylation by ATR at S345, as well as a milder increase in CHK1-phosphorylation upon aphidicolin treatment (Fig. 5F, Supplementary Fig. 10A, D). This suggests that the loss of the 5q arm leads to a reduced CHK1-dependent signaling, which can provide a proliferative advantage in polysomic cells suffering from replication stress. How exactly the loss of chromosome 5q contributes to reduced CHK1 dependent signal is a subject of further investigation.

Strikingly, examination of replication factors and CHK1-dependent signaling in both reverted disomic clones Hdi5 1 and Hdi5 2 uncovered an increase in CHK1 phosphorylation along with a low abundance of MCM7 (Fig. 5F, Supplementary Fig. 10E). While it is possible that additional disadvantageous mutations may have accumulated in the two independently engineered disomic strains, we favor the hypothesis that restoring disomy imposes an additional burden on the previously aneuploid cells. Thus, the removal of a supernumerary chromosome from established trisomies does not necessarily translate to enhanced cellular proliferation. Our data show that arm-level chromosome 5 copy number aberrations have a strong effect on cell proliferation. Strikingly, similar effects can be observed when analyzing the patient survival of TCGA patients. Here, a gain of the p-arm exhibits significantly increased hazard compared to copy number neutral tumors exclusively when there is no coinciding gain of the q-arm (Fig. 5G), corroborating the different contribution of 5q and 5p chromosome arms to malignancy. Together, our analysis provides an example of specific chromosomal aneuploidy that corelates with proliferation and patient prognosis, and provides firs insights into possible molecular mechanisms.

## Discussion

The gain of even a single additional chromosome disrupts cellular homeostasis, exerting adverse effects on most cell types analyzed so far. Whether resulting from spontaneous or induced chromosome missegregation, or through chromosome transfer, the emergence of chromosomal gain in human somatic cells leads to diminished proliferation, pronounced alterations in gene expression, defects in protein homeostasis, compromised DNA replication, accumulation of DNA damage, and the initiation of sterile inflammatory responses, collectively constituting so-called aneuploidy-associated stresses (*2, 41, 42*). These features are mirrored in aneuploid cancers, where replication stress and DNA damage, protein folding defects, and inflammation are frequently observed in tumors (*43–45*). Intriguingly, cancer cells proliferate efficiently despite these challenges, suggesting the possibility that adaptive changes accumulate during tumorigenesis to counteract the negative consequences of aneuploidy. To identify pathways contributing to the adaptation to the deleterious effects of chromosomal gains, we used isogenic human cell lines with an additional chromosome and subjected them to *in vitro* evolution experiments. We observed an improvement in proliferation within these cell lines, and remarkably, this enhancement did not depend on the loss of the extra chromosome (Fig. 1A-C). Instead, the cell lines retained either a part or the entirety of the additional chromosome while accumulating further chromosomal abnormalities. The persistent presence of the supernumerary chromosome provided a unique opportunity for elucidating the physiological changes necessary for adapting to chromosomal gains.

In our quest to detect the factors underlying the enhanced proliferation, we determined the protein abundance in the evolved cell lines and compared it to the corresponding unevolved cells as well as to the parental diploid wild type. This examination pointed out alterations in the expression of genes required for DNA replication and lysosomal functions. Strikingly, the comparison with data provided by TCGA and CPTAC databases showed that the adaptive changes observed in the evolved clones correlate well with the newly established AADEPT score, which quantifies the correlation of gene expression changes observed in primary tumors with the AS (Fig. 2B-D). This correlation was evident at both the gene level, as well as when assessing pathway changes (Fig. 2D, E). Our findings underscore the utility of our model systems in identifying factors relevant to the adaptive responses to aneuploidy in the context of malignant aneuploid tumors.

Among the most striking features we found changes in the abundance of proteins related to DNA replication. While initial chromosome gain leads to reduced expression of proteins required for replication, their abundance increases after *in vitro* evolution (Fig. 2). The same trend is observed in tumor tissues compared to healthy, non-transformed tissues. This is in line with observation in cancers, where high abundance of proteins required for replication correlates often with poor prognosis. Additionally, the replication stress and genomic instability is reduced in evolved in aneuploid. cells (Fig. 3E-H). We conclude that high levels of genomic instability are adversaries of proliferation.

To further identify what molecular mechanisms may contribute to the altered expression of replicative factors and other pro-proliferative proteins after *in vitro* evolution, we identified the most overrepresented proteins after *in vitro* evolution and the responsible transcription factors. Our observations from the *in vitro* evolution revealed high enrichment of proteins associated with M phase, DNA replication, and the mitotic cell cycle, which is consistent with previous findings of factors overexpressed in aggressive tumors (Fig. 2 E) (*46, 47*). We found that consensus targets of transcription factors E2F4 and FOXM1 are strongly enriched in our *in vitro* evolution model, as well as in aneuploid cancers (Fig. 4A). This suggests that to overcome the burdens of aneuploidy, increased expression of specific transcription factors promoting cell-cycle progression is required.

While our comparison with cancer data shows a remarkable similarity among pathways identified in *in vitro* evolution of aneuploid cells and pathways, whose changes during tumorigenesis correlate with aneuploidy (AADEPT), the situation *in vivo* will be certainly more complex. For example, aneuploidy is associated with presence of cytosolic DNA which induces inflammatory pathways, suggesting that aneuploid cells might be targeted by immune cells. We observed reduced expression of several innate immune response genes after *in vitro* evolution (Fig. 4A). Interestingly, aneuploid tumors are largely devoid of immune cells (*48*), suggesting that reduced immunogenicity is associated with aneuploidy and important for propagation of aneuploid cancers.

We were particularly intrigued by the clear association of FOXM1 targets with the adaptation to aneuploidy. FOXM1 is upregulated during early cancer development and exerts multiple tumor-promoting activities through stimulation of cell cycle progression and suppression of senescence (*49, 50*). Moreover, FOXM1 has been found to support proliferation of chromosomally unstable and aging aneuploid cells in both mouse and human (*37, 38*). We found that overexpression of FOXM1 in non-evolved polysomic cells enhanced target gene expression and improved their proliferation, while no positive effect was observed in parental diploid cells (Fig. 4H-J). Our findings underscore the pivotal role of FOXM1 in adaptive responses to aneuploidy. While most polysomic cell lines maintained the extra chromosome, trisomy and tetrasomy of chromosome 5 showed frequent loss of the 5q arm after *in vitro* evolution. Gain of 5p arm and loss of 5q arm are also enriched in tumors (Fig. 5A). We used the recently developed ReDACT-TR approach (*23*) for targeted chromosome arm manipulation and engineered unevolved trisomic cell lines to lose 5q and 5p arm, respectively. We showed the adversary effect of 5q trisomy, while cells that lost the extra copy of 5q, but maintained an extra copy of 5p, proliferated significantly better than the parental trisomy (Fig. 5E). We observed a highly congruent pattern in the survival of TCGA cancer patients, with a significantly increased hazard of 5p gain only with lost or neutral 5q (Fig. 5G). We also obtained two diploid clones, most likely due to induced chromosome loss via CRISPR/Cas9 mediated targeting (*39, 40*). Despite the regained disomy, these cell lines showed strong proliferation defect (Fig. 5E). Our findings correspond with recent analysis of the tumor suppressors and oncogenes contribution to cancer aneuploidies, demonstrating a higher density of oncogenes, as for example TERT and DROSHA, on chromosome 5p, while tumor suppressors like MAP3K1 and MAST4 are more frequently encoded on chromosome 5q (*19, 22*). Recently, a concept of aneuploidy addiction has been demonstrated in cancer cell lines where loss of additional chromosomes compromised cancer-like growth (*23*). Our observation that the loss of an entire extra chromosome 5 from trisomic cell lines reduced proliferation in comparison to the full trisomy, while loss of 5q only and maintaining 5p led to generally improved proliferation, further accentuates the nuanced effects of aneuploidy on cancer cell proliferation.

In summary, we propose that aneuploidy initially impairs cellular fitness through increased proteotoxic stress and expression of lysosomal proteins, as well as increased replication stress and low expression of replication factors, however, cells can adapt to these adverse effects through increased expression of FOXM1 and E2F4 targets or loss of anti-proliferative chromosome segments. One limitation of this study is the limited amount of analyzed polysomic cell lines. However, the cross-comparison of changes identified *in vitro* with pathway regulation in cancer patients showed remarkable convergence. Our approach provides novel insights into molecular mechanisms of cellular adaptations to aneuploidy and may open a new route to potential cancer therapies.

## Material and Methods

### Cell culture and treatment

RPE1 hTERT (referred to as RPE1) and RPE1 hTERT H2B-GFP were a kind gift of Stephen Taylor (University of Manchester, UK). HCT116 H2B-GFP was generated by lipofection (FugeneHD, Roche) of HCT116 (ATCC No. CCL-247) with pBOS-H2B-GFP (BD Pharmingen) according to the protocols from the manufacture. Trisomic and tetrasomic cell lines were generated by microcell-mediated chromosome transfer as described in (*3*) (see below). All cell lines were cultured at 37 °C with 5% CO_2_ atmosphere in Dulbecco’s Modified Eagle Medium (DMEM) (Gibco™) containing 10% fetal bovine serum (FBS) (Gibco™) and 1% Penicillin/Streptomycin (PenStrep) (Gibco™).

### Microcell-mediated chromosome transfer

Mouse A9 donor cells containing a single human chromosome and tagged with a drug-resistant gene were treated with colchicine for over 48 hours to induce micronucleation. The microcells with micronuclei containing individual chromosomes were isolated through centrifugation with cytochalasin B. The filtered micronuclei were then treated with phytohemagglutinin-P (PHA-P) to allow adherence to the recipient cells. Polyethylene glycol (PEG) was used to fuse the micronuclei to the recipient cells. Cells with additional chromosomes were selected by antibiotic selection respective to the drug-resistant gene. For further details, see (*3, 25*).

### Proliferation assay

RPE1 and HCT116-derived cells were seeded in triplicates into a 96-well plate (RPE1: 1.5×10 ^3^ cells/well; HCT116: 6×10^3^ cells/well). The CellTiter-Glo® Luminescent Cell Viability Assay by Promega was used to assess the proliferation. 24-hour intervals were chosen for measurements. CellTiter Glo reagent was added to each well according to the manufacturer’s instructions. The luminescence was measured using the Promega GloMax Explorer plate reader. Values for each cell line were standardized to day 0 of the respective sample.

### Clonogenic assay

500 cells/well were seeded to 6 well plates and incubated for 10 days at 37°C in 5% CO_2_ incubator. Cells were then washed with PBS, fixed and stained using 20% methanol and 0.5% crystal violet solution for 20 minutes at room temperature. To remove the staining, cells were washed once using PBS and twice using water. After drying, the plates were imaged and colonies were analyzed using Fiji (*51*).

### Senescence and Viability assay

The proportion of dead and senescent cells was determined by flow cytometry of CV450-stained cells and the fluorescent SA-β-Gal substrate DDAO-Galactoside, respectively. The FlowJo™ Software (v10.8, BD Life Sciences) was utilized to analyze the obtained data.

### Metaphase spreads

At a confluency of 70–80%, cells were treated with 400 ng/ml colchicine for 6 hours. Cells were collected by trypsinization and centrifuged at 1500 rpm for 10 minutes. Cell pellets were resuspended in 75 mM KCl and incubated for 10 minutes at 37 °C. Fixation was performed with 3:1 methanol/acetic acid. Fixed samples were dropped onto glass slides. Vectashield Mounting Medium with DAPI was used to visualize the DNA.

### Fluorescence *in situ* hybridization

At a confluency of 70–80%, cells were treated with 400 ng/ml colchicine for 6 hours. Cells were collected by trypsinization and centrifuged at 1500 rpm for 10 minutes. Cell pellets were resuspended in 75 mM KCl and incubated for 10 minutes at 37 °C. Fixation was performed with 3:1 methanol/acetic acid. Fixed samples were dropped onto glass slides. Vectashield Mounting Medium with DAPI was used to visualize the DNA.

### Analysis of DNA replication by EdU incorporation

EdU (10 μM) was added to the medium for 30 minutes at 37 °C. After the incorporation, the medium containing EdU was discarded, and the cells were harvested via trypsinization (5 minutes) and collected into 15 ml falcon tubes. After centrifugation for 3 minutes at 1300 rpm (Rotina 420R), the cell pellet was washed with PBS and centrifuged again (3 minutes, 1300 rpm, Rotina 420R). The supernatant was discarded, and the cells were permeabilized for 15 minutes with Fix/Perm solution (Thermo Fisher Scientific) according to the manufacturer’s instructions. After permeabilization, 1 ml PBS was added, and the samples were then centrifuged for 3 minutes at 1300 rpm (Rotina 420R). The supernatant was discarded, and the cell pellets were resuspended in 100μl Perm wash (Thermo Fisher Scientific). Subsequently, 500 μl Click-iT Reaction Mix (1μM Eterneon Red (baseclick GmbH), 6.6% (v/v) 1.5 M Tris (pH 8.8), 500 μM CuSO4, 100 mM Ascorbic Acid in PBS) was added to the samples for 20 minutes at room temperature in the dark. After the incubation, the samples were washed 3 times with 500 μl Perm wash (centrifugation for 2 minutes at 1600 rpm (Eppendorf centrifuge 5415R)). The samples for the pRPA2 (S33) analysis were additionally incubated in 100 μl 1:500 pRPA2 (S33) antibody and subsequently in 100 μl 1:500 Alexa Fluor 594-Goat anti-rabbit for 30 minutes in the dark, each. Finally, the DNA was stained by incubating the cells in 300 – 500 μl (volume depending on the pellet size) PBS containing 1 μg/ml DAPI and 10 μg/ml RNase. The cells were analyzed by flow cytometry (see below).

### Flow Cytometry analysis of EdU-incorporated cells

The flow cytometry was performed with an Attune™ NxT Flow Cytometer (Invitrogen) following the manufacturer’s instructions. DAPI and EdU-bound Eterneon-Red were excited by a 405 nm and a 638 nm laser, respectively. The emission from the excited DAPI was collected with a 440/50 filter. The light emitted by the excited Eterneon-Red was collected with a 670/14 filter. The FlowJo™ Software (v10.8, BD Life Sciences) was utilized to analyze the obtained data. First, the population of cells was gated by excluding particles based on the pulse area of the Side and Forward Scatter. Subsequently, the population of single cells was gated from the population of cells by pulse height versus pulse area of Forward Scatter. Finally, the pulse area of the Eterneon-Red and the DAPI emissions were plotted against one another to assign the single cell sub-populations based on their DAPI and Eterneon-Red emission displaying the DNA content and the EdU incorporation, respectively.

### Protein isolation

Cells were either seeded into 6 cm dishes (1×10^6^ cells per dish) or into 10 cm dishes (2.2×10^6^ cells per dish). As a positive control, cells were treated with 1 μM doxorubicin or other drugs (as specified in the figures) and incubated overnight to trigger DNA damage response. For harvesting, the cells were detached from the dish via trypsinization (5 minutes) and collected into 15 ml falcon tubes. After centrifugation for 3 minutes at 1300 rpm (Rotina 420R), the cell pellet was washed with PBS and centrifuged again (3 minutes, 1300 rpm, Rotina 420R). The supernatant was discarded, and the cell pellet was resuspended in 50-200 μl RIPA buffer (volume depending on the pellet size) supplemented with protease (Roche Diagnostics GmbH) and phosphatase inhibitors (Roche Diagnostics GmbH). The samples were sonicated on ice for 12 to 15 minutes and then centrifuged for 10 minutes at 13200 rpm and 4 °C (Eppendorf centrifuge 5415R) to remove the cell debris. The pellets were thrown, and the protein concentrations of the supernatants (whole-cell protein lysates) were determined in a Bradford protein assay. The protein concentrations were then adjusted to either 1 μg/μl or 10 μg/μl in 1x Laemmli buffer. After the samples were boiled for 5 to 10 minutes at 95 °C they were stored at 20 °C until further usage.

### SDS-PAGE and immunoblotting

Depending on the well size 10 to 15 μg of total protein was loaded on the gel per well. The Precision Plus Protein All Blue Standard (BioRad, Hercules, USA) or the Color Prestained Protein Standard, Broad Range (NEB) was used as a molecular weight marker. For stacking, gels were run at a constant voltage of 85 to 100 V for approximately 30 minutes. After that, the voltage was increased to 120-150 V for separation. Subsequently, either a wet transfer or a semidry transfer was used to transfer the proteins to a nitrocellulose blotting membrane. For the semidry transfer, Trans-Blot® Turbo™ (BioRad Laboratories, Hercules, USA) and Bjerrum-Schaefer-Nielsen buffer were used. The wet transfer was either done overnight at a constant current of 0.16 A or for 1 hour at a constant voltage of 100 V (both at 4 °C). The transfer was verified by staining the membranes with Ponceau solution for 5 minutes. Nonspecific binding sites were blocked by incubating the membranes in 5% milk in TBS-T for 1 hour at room temperature. After blocking, the incubation with the primary antibodies diluted in 5% milk or 5% BSA in TBS-T was done overnight at 4 °C. One day later, the membranes were washed 3 times with TBS-T for 5 minutes before the corresponding secondary antibodies (conjugated with HRP) diluted in 5% milk in TBS-T were added. This was incubated for at least 1 hour at room temperature. Unbound secondary antibodies were removed by washing again 3 times for 5 minutes with TBS T. Finally, the protein signals were detected by using the Clarity™ Western ECL substrate solution (BioRad) and the Azure c300 system (Azure Biosystems, Dublin, USA). For quantification, the signal intensities were measured using the software ImageJ (v. 1.52i). All used antibodies are listed in the table below.

### Flow cytometry of CHK1 phosphorylation

Cells were seeded to 6 well plates and incubated (37 °C, 5 % CO_2_) for 24 hours. Then, 200 nM aphidicolin was added to the cells and incubated again for 24 hours. Cells were harvested using trypsin and washed using PBS. Fixation was performed using 70% EtOH for at least 30 minutes at 4 °C, followed by permeabilization for 15 minutes (0.25% Triton X-100 in PBS). Subsequently, the cells were incubated in PBS+1% BSA for 30 minutes at RT. After blocking, the cells were incubated with primary antibody solution (1% BSA in PBS; pCHK1 primary AB 1:1000) at 4 °C overnight. Upon removal of the primary antibody solution, the cells were incubated with secondary antibody solution (1% BSA in PBS; AlexaFluor647 anti-rabbit 1:300) for 30 minutes at room temperature. Finally, staining solution containing DAPI (0.1 μg/ml) and RNase A (0.01 mg/mL) in PBS was used to stain the DNA for 5 min. Fluorescence intensities were measured using Attune™ NxT Flow Cytometer (Invitrogen) and analysis was performed using FlowJo™ Software (v10.8, BD Life Sciences).

### Competition assay

HCT116 or Htr5 cells overexpressing FOXM1, and control cells were transduced with the pMMLV-EBFP-P2A-Puro or pMMLV-mRFP1-P2A-Puro, respectively, or vice versa. The cells were mixed in a 1:1 ratio and the proportion of cell fractions expressing RFP or BFP was quantified by flow cytometry after 1 day, 4 days, 7 days, and 10 days. For each measurement, 10.000 cells were harvested per mix and the cells were resuspended in PBS with 2 mM EDTA and the fluorescence ratio was measured with the SH800S Cell Sorter (Sony Biotechnology, Weybridge, UK). The rest of the cell mix was passaged with each measurement. The ratio between RFP- and BFP-positive cells was analyzed using FlowJo™ Software (v10.8, BD Life Sciences).

### Immunofluorescence staining

Cells were seeded in black glass bottom 96-well plates (Thermo Fisher ScientificTM) and cultured in DMEM (with FBS and PenStrep) to a confluence of 70–80%. Fixation was performed for 12 min at RT with freshly prepared 3:1 methanol:acetic acid or with 3% formaldehyde. As permeabilization buffer, 0.5% Triton X-100 (Carl Roth) in PBS was used for a 5-minute incubation at RT. Subsequently, the cells were blocked in 0.1% BSA for 30 minutes at RT. Primary antibody incubation was performed overnight at 4 °C. Subsequent washing was done 3x with PBS. The secondary antibody incubation was performed for 1 hour at RT. The DNA was counterstained with SytoxGreen (167 nM) or 1.0 mg/ml DAPI both containing RNase A (0.01 mg/ml) and imaged as described below. Plates were stored in PBS with NaN_3_.

### Lysotracker staining

0.2×10^6^ cells/well were seeded in 96-well plates. The following day, the cells were washed with PBS and replaced with medium containing 500 nM Lysotracker for 4 hours at RT. Subsequently, the cells were washed with PBS and fixed in 3:1 methanol:acetic acid. After repeated rinsing with PBS-T, nuclei were counterstained with Sytox Green (167 nM) containing RNase A (0.01 mg/ml). Plates were sealed with parafilm following the addition of PBS containing sodium azide (1 mM).

### DNA staining for microscopy

2.5×10^4^ cells/well were seeded in 96-well plates. The positive control was treated with 0.2 μM aphidicolin and 0.73 mM caffeine for 24 hours. Cells were fixed in 3:1 methanol:acetic acid. Subsequently, the cells were washed with PBS-T. The DNA was stained with Sytox Green (167 nM) containing RNase A (0.01 mg/ml). The plates are sealed with parafilm after the addition of PBS containing sodium azide (1 mM).

### DNA staining for microscopy

2.5×10^4^ cells/well were seeded in 96-well plates. The positive control was treated with 0.2 μM aphidicolin and 0.73 mM caffeine for 24 hours. Cells were fixed in 3:1 methanol:acetic acid. Subsequently, the cells were washed with PBS-T. The DNA was stained with Sytox Green (167 nM) containing RNase A (0.01 mg/ml). The plates are sealed with parafilm after the addition of PBS containing sodium azide (1 mM).

### DNA combing

2×10^5^ cells were pulse-labelled for 30 minutes with 10 mM CldU 5-chloro-2’-deoxyuridine (CIdU) followed by 30 minutes of 10 mM CldU 5-iodo-2’deoxyuridine (IdU) according to the Genomic Vision Replication Combing Assay (RCA) Protocol (Paris, France). DNA was extracted using the Genomic Vision Fiber-Prep Kit for DNA extraction and Fibers were stretched following manufacturer’s instruction using the Genomic Vision RCA Protocol. A Genomic Vision FiberComb device with a stretching factor (SF) of 2 was used. Antibody staining was performed according to the RCA Protocol.

### Microscopy

Microscopy was performed using a semi-automated Zeiss AxioObserver Z1 (Oberkochen, Germany) equipped with an ASI MS-2000 stage (Applied Scientific Instrumentation, Eugene, USA), a CSU-X1 spinning disk confocal head (Yokogawa) and a Laser Stack with selectable lasers (Intelligent Imaging Innovations, Denver, USA) and the Cool-Snap HQ camera (Roper Scientific). 40x air or 63x oil objectives were used under the control of the software Slidebook6 (Intelligent Imaging Innovations, Denver, CO). Foci analysis was performed automatized using Cell Profiler (v4.2.1).

### Lentiviral infection

HEK 293T cells were transfected with the packaging plasmids pHDM-Hgpm2, pHDM-Tat1b, pHDM-VSVG, and pRC-CMV-Rev1b (kind gift from Alejandro Balazs), and pLVX–Tight-Puro–FoxM1-dNdK (kind gift from Elsa Logarinho), pMMLV-EBFP-P2A-Puro, or pMMLV-mRFP1-P2A-Puro (both were a kind gift from Jason Sheltzer (Addgene plasmid #160227; http://n2t.net/addgene:160227; RRID:Addgene 160227) (Addgene plasmid #160228; http://n2t.net/addgene:160228 ; RRID:Addgene 160228)) by using Lipofectamine™ 2000 (Life Technologies, Thermo Scientific, CA, USA) according to the manufacturer’s guidelines. The recipient cells were infected 48 hours post-transfection of HEK 293 cells with the virus supernatant in the presence of 8 μg/ml polybrene according to the manufacturer’s guidelines (Sigma-Aldrich, MO, USA). The cells transduced with the pLVX-Tight-Puro-FoxM1-dNdK were selected in 2 μg/ml puromycin (Gibco, Thermo Fisher Scientific, CA, USA).

### Induced chromosome arm-loss

We used the previously described REDACT-TR system (*23*). For target site prediction, we used the E-CRISP gRNA design tool (*52*), using reference genome hg38. The corresponding gRNA sequences were cloned into the pX330-U6-Chimeric BB-CBh-hSpCas9 backbone (gift from Feng Zhang; Addgene plasmid #42230) using BbsI (NEB) restriction sites. To introduce a selection marker via NHEJ, a fragment cleaved from EF1a Puro Telo v1 (gift from Jason Sheltzer; Addgene plasmid #195138), which encodes a puromycin resistance cassette fused to a telomeric sequence, was used. A fragment produced by KpnI (NEB) and BstZ17l (NEB) digest was extracted from an agarose gel and co-transfected with the pX330 plasmid into the Htr5 cell line using the PEI transfection method. Two days after transfection, 1 μg/ml puromycin was added for 48 hours to select the transfected cells. Single colonies were picked two weeks later. The resulting cell lines were screened by FISH (Metasystems) and validated by sequencing.

### Statistical analysis

The R software (version 4.2.1) (*53*) was used to perform growth curve quantification as well as multi-omic data analysis of polysomic model cells lines as well as TCGA and CPTAC patient samples. For visualization the R packages ggplot2 (v3.4.4) (*54*), gghx4 (v0.2.6), ggpubr (v0.6.0) and cowplot (v1.1.1) were used. For all other experimental data, analysis was done in GraphPad Prism 9.

### Growth curve quantification

The raw luminescence values in the results of the CellTiter-Glo® assays were normalized by subtracting the RLU measured at the initial time points. Then, a linear spline model was fit through the normalized luminescence values separately for each cell line and experimental run using the R package lspline (v1.0). Residual bootstrapping of the fitted models was performed to create R = 10000 bootstrap samples using the Boot function of the car (v3.0) (*55*) package. For each bootstrap sample, the area under the curve (AUC) of the linear spline was calculated by summation of the trapezoid areas between the horizontal axis and the line between two adjacent time points. The resulting AUCs were divided by the respective estimates of the wild-type cell lines at passage 0 and averaged over the different experimental runs. Finally, empirical 95% confidence intervals for the average AUC fold change were calculated using the 2.5% and 97.5% quantiles of the bootstrap samples.

### Genomic analysis

Genome-wide comparative genomic hybridization of cell lines (Supplementary Table 2) was carried out by IMGM laboratories (Martinsried, Germany). Agilent Human Genome CGH Microarrays (4×44K format) and SurePrint G3 Human CGH Microarray (4×180K format) were used in combination with a Two-Color based hybridization protocol with a commercially available reference gDNA. Signals on the microarrays were extracted using the Agilent DNA Microarray Scanner. The Agilent Genomic Workbench 7.0 was used on the raw feature extraction data to apply probe filtering, log2 transformation, GC correction and re-centralization to obtain log2 copy number ratios relative to the reference gDNA. Further details are provided with the uploaded data files (see Data and Code availability).

Low-coverage whole genome sequencing (lcWGS) of cell lines was performed at the NIG Integrative Genomics Core Unit (Göttingen, Germany) in two batches (see Supplementary Table 2). The genomic DNA was extracted in-house. Cells were collected with trypsin and centrifuged. The pellet was washed twice with PBS. DNA isolation was performed using the Qiagen DNA easy Blood and Tissue Kit (Hilden, Germany) following manufacturer’s instructions. gDNA from each cell line was used for library preparation and paired-end sequencing (151 bp read length) to an average depth of >1x on an Illumina HiSeq4000 machine. For data from batch 1, raw reads were aligned to the human reference genome (GRCh37/hg19) using the Illumina DRAGEN Bio-IT Platform (Host Software version 05.021.609.3.9.5 and Bio-IT Processor Version 0×04261818) to generate BAM files, and for batch 2, alignment was done using the BWA-MEM algorithm (v0.7.17) (*56*) and conversion to BAM format using Samtools htslib (v1.9) (*57*).

Mini-bulk sequencing was performed at the ERIBA University Medical Center (Groningen, Netherlands) and cell lines were sent there for preparation and DNA extraction (see Supplementary Table 2). For sequencing, cells were suspended in media, washed, and pelleted. To generate nuclei, cells were resuspended in cell lysis and staining buffer (100 mM Tris-HCl pH 7.4, 154 mM NaCl, 1 mM CaCl_2_, 500 μM MgCl_2_, 0.2% BSA, 0.1% NP-40, 10 μg/mL Hoechst 33358, 2 μg/mL propidium iodide in ultra-pure water). Resulting cell nuclei were gated for G1 phase (as determined by Hoechst and propidium iodide staining) and sorted into wells of 96 wells plates containing freezing buffer on a MoFlo Astrios cell sorter (Beckman Coulter). 30 nuclei of each were deposited per well. Plates containing nuclei and freezing buffer were stored at -80°C until further processing. Automated library preparation was then performed as previously described (*58*). Libraries were sequenced on a NextSeq 2000 machine (Illumina; up to 77 cycles – dual-index, single end), and aligned to the human reference genome (GRCh38/hg38) using Bowtie2 (v2.2.4 or v2.3.4.1) (*59*). Duplicate reads were marked with BamUtil (v1.0.3) (*60*) or Samtools markdup (v1.9) (*57*).

### Extraction and segmentation of genomic copy number ratios

For the aligned reads from lcWGS and minibulk sequencing, the tools from the HMM Copy Utils repository (https://github.com/shah-compbio/hmmcopy_utils) were used for extracting read counts from the BAM files in 50 kb (lcWGS) and 0.75 Mb (minibulk) bins as well as GC and mappability statistics from the reference genomes. The R package HMMcopy (v1.38) was then used to estimate log2 copy number ratios controlling for mappablilty and GC contentEstimated copy number ratios from aCGH, lcWGS and minibulk sequencing were then normalized by subtracting the values of the respective unevolved wild type. The normalized log2 copy number ratios were then grouped based on genomic location using circular binary segmentation as implemented in the R package DNAcopy (v1.70) and average and median copy number ratios were calculated per segment.

### Estimation of total relative DNA

The amount of total relative autosomal DNA was estimated heuristically as: 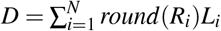, where *N* is the number of distinct segments of autosomal chromosomes identified by CBS, *L*_*i*_ is the length of segment *i* in bp and Ri is the approximate difference in total copy number (which is being rounded up to the nearest digit), calculated as: 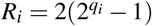. Here *qi* is the normalized, log2 copy number ratio of segment *i* as calculated above. Assuming that the reference copy number is approximately the same for each cell line of interest and the unevolved, wild type, it follows, that: 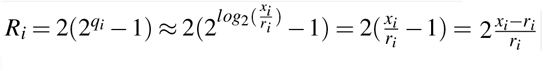 where *xi* corresponds to the copy number of segment *i* in the cell line of interest and *ri* to the corresponding value in the wild-type control. Now, reasoning that the parental cell lines are diploid, expect for a few gains that do not change with the addition of extra chromosomes or in-vitro evolution (Fig. 1C), it is assumed that:

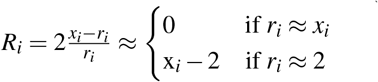

Thus, under the above assumptions, *Ri* is indeed approximately the difference in segment copy number when being rounded to the nearest integer, and *D* corresponds to the total amount of autosomal DNA relative to wild type.

### Correlation analysis

Calculation and statistical testing of Pearson and Spearman correlation coefficients was performed using the cor.test function of the base R stats package with default parameters.

### Sample preparation for tandem mass tag (TMT)-based quantitative proteomics

Cultured cells (1×10^6^ cells) were harvested by trypsinization, centrifuged at 1,200 rpm for 3 minutes and washed 1x with 1xPBS. Pellets were snap-frozen in liquid nitrogen and stored at -80°C. Sample preparation and labeling of the peptides with TMT isobaric mass tags was performed as manufacturer’s instructions (Thermo Fisher Scientific). In brief, cells were lysed with provided lysis buffer and DNA was sheared by sonication in a water bath for 20 cycles (30 seconds on / 30 seconds off). After centrifugation at 12,000 x g for 10 minutes at 4 °C, protein concentration was determined using the bicinchoninic acid protein assay kit (Thermo Fisher Scientific) (PierceTM BCA Protein Assay Kit, Thermo, #23225). 50 μg of protein per condition was reduced with 5 mM Tris(2-carboxyethyl)phosphin -hydrochlorid T (TCEP) for 1 hour at 55 °C and subsequently alkylated with 10 mM iodacetamide for 30 min at 25 °C protected from light. By adding 6 volumes of acetone, proteins were allowed to precipitate overnight at -20 °C. Precipitated samples were resuspended in 100 mM TEAB and digested by incubation with 1.5 μg proteomics-grade trypsin (Sigma-Aldrich #T6567) at 37 °C overnight. 25 μg of peptides per condition were labelled with isobaric tags by incubation for 1 hour at RT (TMT 6-plex and 10-plex, Thermo Fisher Scientific, TMTsix/tenplex™ Isobaric Mass Tagging Kit, #90061, #90406). For 10-plex and 6-plex analysis cell line samples were individually tagged and for each cell line three biological replicates were labelled separately. The reaction was quenched by adding 5% hydroxylamine to the samples and incubating the samples for 15 min at RT. After combining all individually labelled samples per replicate, TMT-labeled peptides were fractionated into 8 fractions using the high pH reverse-phase peptide fractionation kit according to manufacturer’s instructions (Thermo Fisher Scientific) (Pierce High pH Reversed-Phase Peptide Fractionation Kit, Thermo Fisher Scientific, #84868) and subsequently dried using vacuum centrifugation.

### Liquid chromatography-tandem mass spectrometry

TMT-labeled peptides were dissolved in 0.5% acetic acid supplemented with 1% of 2% acetonitrile / 0.1% TFA pH2 and analyzed by liquid chromatography-tandem mass spectrometry (LC-MS/MS) with a system of an EASY nano-LC 1200TM system (Thermo Fisher Scientific) and a Q Exactive HF (Thermo Fisher Scientific) connected through a Nanospray Flex Ion Source (Thermo Fisher Scientific). 3 μl of each fraction was separated on a 40 cm heated reversed phase HPLC column (75 μm inner diameter with PicoTip EmitterTM, Nex Objective) in-house packed with 1.9 μm C18 beads (ReproSil-Pur 120 C18-AQ, Dr. Maisch). Peptides were loaded in 5% buffer A (0.5% aequeous formic acid) and eluted with a 3 hour-gradient (5-95% buffer B (80% acetonitrile, 0.5% formic acid) at a constant flow rate of 0.25 μl/ml). Mass spectra were obtained in a data-dependent mode. In brief, each full scan (mass range 375-1400 m/z, resolution of 60,000 at m/z of 200, maximum injection time 80 milliseconds, ion target of 3E6) was followed by high-energy collision dissociation based fragmentation (HCD) of the 15 most abundant isotope patterns with a charge state between 2 and 7 (normalized collision energy of 32, an isolation window of 0.7 m/z, resolution of 30,000 at m/z of 200, maximum injection time 100 milliseconds, AGC target value of 1E5, fixed first mass of 100 m/z and dynamic exclusion set to 30 seconds). Raw MS data were processed with the MaxQuant software (v1.6.3.3). All data was mapped to the human reference proteome database (UniProt: UP000005640) with peptides with an FDR ≤ 1%. The TMT mass spectrometry data set three (See Supplementary Table 2) was acquired from the Proteomics Core Facility (Basel, Switzerland). Sample preparation and tandem mass tag labelling, mass spectrometric analysis as well as database searching and protein quantification were all performed as previously described (*11*). The acquired TMT reporter ion intensities from peptides, that were identified in all replicates, were summed over corresponding protein accession numbers with missing values replaced by half of the lowest measured value and values over technical replicates being averaged.

### Protein and gene annotation

Mapping between UniProt protein identifiers and NCBI and Ensembl gene identifiers as well as chromosome, band and genomic location information was done using the biomaRt (v2.52.0) (*61*) R package.

### Differential protein abundance analysis

Identified protein groups were filtered to remove contaminants, reverse hits and proteins identified by site only. Protein groups with missing intensities were removed leaving sets of 6300, 5920 and 6140 robustly identified protein groups in total for TMT experiment one to three respectively (See Supplementary Table 2). The remaining TMT reporter intensities were log2-transformed, cleaned for batch effects using the R package limma (v3.52.4) (*62*) and normalized using variance stabilization normalization between samples as implemented in the vsn package (v3.64.0) (*63*).

Pair-comparisons of protein intensities between cell lines were carried out using weighted linear regression models with observation-specific precision weights from the voom method and empirical Bayes moderated t-statistics from the eBayes method as implemented in limma. The resulting p-values were FDR-adjusted according to the Benjamini-Hochberg method. The analysis results from the three TMT experiments were joined by the proteins corresponding Ensemble gene identifiers.

### Estimation of dosage compensation

For a comparative view with the segmented, genomic, log2 copy number ratios, the log2 protein abundance fold changes were grouped based on the genomic location of their corresponding genes using circular binary segmentation as implemented in the R package DNAcopy (v1.70). Chromosome arm-level dosage compensation was calculated as the difference between average genomic log2 copy number ratios and the average log2 abundance fold change of proteins originating from the respective arm, all relative to the unevolved parental cell line.

### Preparation of multi-omics patient data sets

Cancer patient-derived multi-omics and clinical data of TCGA was obtained as collected by Pan-Cancer Atlas project publications (*33*). Aneuploidy scores and arm-level copy number gain and loss calls as well as whole genome doubling status were taken as calculated by (*21*) using the ABSOLUTE algorithm (*64*). Merged and batch-corrected pan-cancer RSEM RNAseq data was taken from (*33*) and processed like in the source paper, filtering for genes with valid expression values in at least 60% of samples and applying upper-quartile normalization and log2 transformation. To make the tumor data more comparable to the situation in our model cell lines, patient with tumors that have undergone whole genome doubling were excluded. Filtering further for patients with tumors that have both genomic and transcriptomic data as well as matching data of normal tissue, a set of 467 patients with 22 different cancer types was derived. Finally, the processed gene expression values of normal tissues were subtracted from the corresponding primary tumor data, resulting in a set of 14282 genes with normalized expression values (Supplementary Fig. 5A, Supplemental Table 4).

Corresponding data from the CPTAC was obtained from the Pan Cancer Analysis resources (*29*) Whole exome sequencing copy number variation data as processed by the Washington University team’s pipeline was used to obtain segmented log2 copy number ratios as well as ploidy calls derived from ABSOLUTE. Here, we calculated the aneuploidy score as in (*21*) by determining the number of autosomal chromosome arms with either a copy number gain or loss covering more than 75% of its length. As recommended in (*29*) a chromosomal segment was determined to be gained or lost when the corresponding absolute, log2 copy number ratio exceeded 0.2. To likely filter out patients with tumors that have undergone whole genome doubling, we set a cut-off of 2.5 for the ABSOLUTE-derived ploidy. This resulted in aneuploidy scores with a comparable distribution as in TCGA patients (Supplementary Fig. 5A, Supplementary Fig. 5B). Pan-Cancer proteome data as processed by the University of Michigan team’s pipeline, and then post-processed by the Baylor College of Medicine’s pipeline was filtered to only include patients with corresponding data between tumor and normal tissue. Consequently, the processed log2, relative protein abundance values of normal tissues were subtracted from the tumor data. Then, filtering further for patients with matching genomic and proteomic data and merging all 6 remaining cancer types, we derived a set of 347 patients with normalized protein values mapping to 8404 different genes (Supplementary Fig. 5B, Supplemental Table 4).

### AADEPT score calculation

Aneuploidy-associated differential expression in tumorigenesis (AADEPT) scores were calculated by determining Spearman rank correlation coefficients between the aneuploidy score in the patient tumors and the mRNA gene expression (TCGA) and protein abundance (CPTAC) values contrasted with corresponding normal tissue samples. Subsequently, the correlation coefficients were tested for statistical significance using the cor.test function of the base R stats package.

### Enrichment analysis

For enrichment analysis the previously developed 2D enrichment method developed by (*65*) was extended to be applicable to any number of dimensions. Accordingly, log2 protein abundance fold changes of cell line pair comparisons or protein AADEPT scores from CPTAC or TCGA tumors were first ranked and scaled from -1 (lowest) to 1 (highest). Then, for one-dimensional enrichment, the ranked values of proteins belonging to a set were tested for significant mean difference when compared to the respective background distribution of all other quantified proteins using univariate ANOVA as implemented in the stats package anova function. The observed difference in average protein ranks constitutes the enrichment score. Likewise, for multidimensional enrichment, protein fold change and/or AADEPT score ranks from multiple comparisons were taken as the input for multivariate ANOVA as implemented in the stats package’s manova function to test for overall difference in means relative to the background. This way, gene set enrichment can be tested globally for a set of comparisons of arbitrary dimension. Lastly, FDR-adjustment for simultaneous testing of KEGG pathways or hallmark gene sets was done using the Benjamini-Hochberg method.

### Hierarchal clustering

Heatmap clustering of proteins and protein sets was performed using agglomerative clustering as implemented in the R stats package function hclust with default parameters.

### Protein relevance score

The relevance of protein abundance change patterns in our evolving, polysomic model cell lines was accessed by calculating protein-wise relevance scores (Supplementary Fig. 5C). These were derived as followed: First, the abundance fold changes of a protein in all pairs of cell line comparisons were ranked from -1 (most negative) to 1 (most positive). Then, a protein’s fold change ranks were grouped based on the type of cell line comparison, being either between an unevolved polysomic cell line and its parental wild type (group 1) or between an evolved cell line and its unevolved counterpart. The latter group was further divided into polysomic (group 2) and disomic control evolutions (group 3). Next, the sum of fold change ranks in group 1 and group 2 were added to one another. To account for changes in protein abundance that could be largely due to the effects of prolonged passaging, the maximum fold change rank in group 3 with equal sign to the sum from group 2 (or 0 if there was no such fold change) was subtracted from the sum prior to taking its absolute value. Only those proteins identified in all proteomics data sets were considered for scoring. Finally, empirical p-values for each proteins observed relevance score was calculated from 100000 bootstrap samples of relevance scores derived from random permutation of proteins.

### Over-representation analysi

Hypergeometric tests were performed to assess the statistical significance of the overlap between the sets of differentially abundant proteins and the different molecular signature sets given all identified proteins as a background. FDR-adjustment of p-values was done according to Benjamini-Hochberg within signature set categories. The strength of the over-representation was calculated as the ratio between the observed and expected overlap, namely: 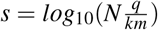 with *q* being the observed overlap, *k* the number of differentially abundant proteins, *m* the size of the signature set and *N* the number of measured proteins among the generated proteomic data sets (N = 7310).

### PEI-transfection

Cells were cultured in 6-well plats to 70% confluency. 2 μg per plasmid/fragment were mixed with sterile-filtered 6 μl PEI-transfection reagent (1 mg/ml) in 700 μl serum-free medium and incubated for 30 minutes at room temperature. The mixture was then added drop-wise to the cells and incubated at 37 °C. 6 hours after transfection cells were washed to remove PEI. Antibiotic selection started 48 hours after transfection.

### Patient survival analysis

The analysis of overall survival of patients included in the TCGA was performed using Cox’s proportional hazards model as implemented in the survival R package (v3.3-1) (*66*) function coxph. Patients were stratified based on the cooccurrence of chromosome 5 arm-level gain or loss events, taking patients with copy number neutral chromosome 5 as the reference group. Only strata with N ≥ 50 were considered. Then, the model was fit using overall survival events and days as the outcome and the strata as well as patient age at initial diagnosis as covariates. Wald test derived p-values were taken as indicators of the statistical significance of each strata’s hazard ratios. The predicted survival curves for each stratum and for a patient of average age (60.7 years) were visualized using the survminer R package (v0.4.9).

## Data availability

### Datasets produced in this study are available in the following databases

- Genomic data from aCGH experiments: GEO GSE254936
- Aligned reads from lcWGS and minibulk sequencing: SRA under BioProject PRJNA1073693

All mass spectrometry proteomics data are available upon reasonable request.

### Downloaded data was accessed through the following repositories

- TCGA data: [PanCanAtlas project page](https://gdc.cancer.gov/about-data/publications/PanCan-CellOfOrigin) of (*33*)
- CPTAC data: [Pan-Cancer Analysis Page](https://proteomic.datacommons.cancer.gov/pdc/cptac-pancancer)
- KEGG pathway and Hallmark gene sets: [Molecular signature data base](https://www.gsea-msigdb.org/gsea/downloads_archive.jsp) (version 2023.1 Mar 2023) (*34*)
- ENCODE and CHEA consensus transcription factor target sets: [EnrichR Gene-set Library](https://maayanlab.cloud/Enrichr/#libraries) (*67*)

Scripts used to analyze the data and generate the figures are available upon request.

## Acknowledgments

The initial observation of the adaptation and improved proliferation of polysomic cells after prolonged passaging was made by Silvia Stingele and Sara Schunter. We thank the students of the Master Program Molecular Cell Biology at the RPTU Kaiserslautern-Landau for their help with the experiments. We thank Elsa Logarinho for providing the FOXM1 overexpressing construct, and Jason Sheltzer for providing the ReDACT-TR and competition assay plasmids (via Addgene). This project was funded by the FOR2800 Chromosome Instability: “Cross-talk of DNA replication stress and mitotic dysfunction” (DFG) to ZS and MR.

## Author Contributions

Conceptualization, Z.S.; Methodology, Z.S., J-E.B., and M.R.; Investigation, K.K., S.R., K.B., L.J., A.W., and M.R.; Writing – Original Draft Z.S.; Writing – Review & Editing, all authors; Computational analysis, Visualization, Software, Formal analysis, and Data curation, J-E.B.; Funding Acquisition, Z.S. and M.R.; Supervision, Z.S. and K.K.

## Supplementary Figures

**Figure S1.**
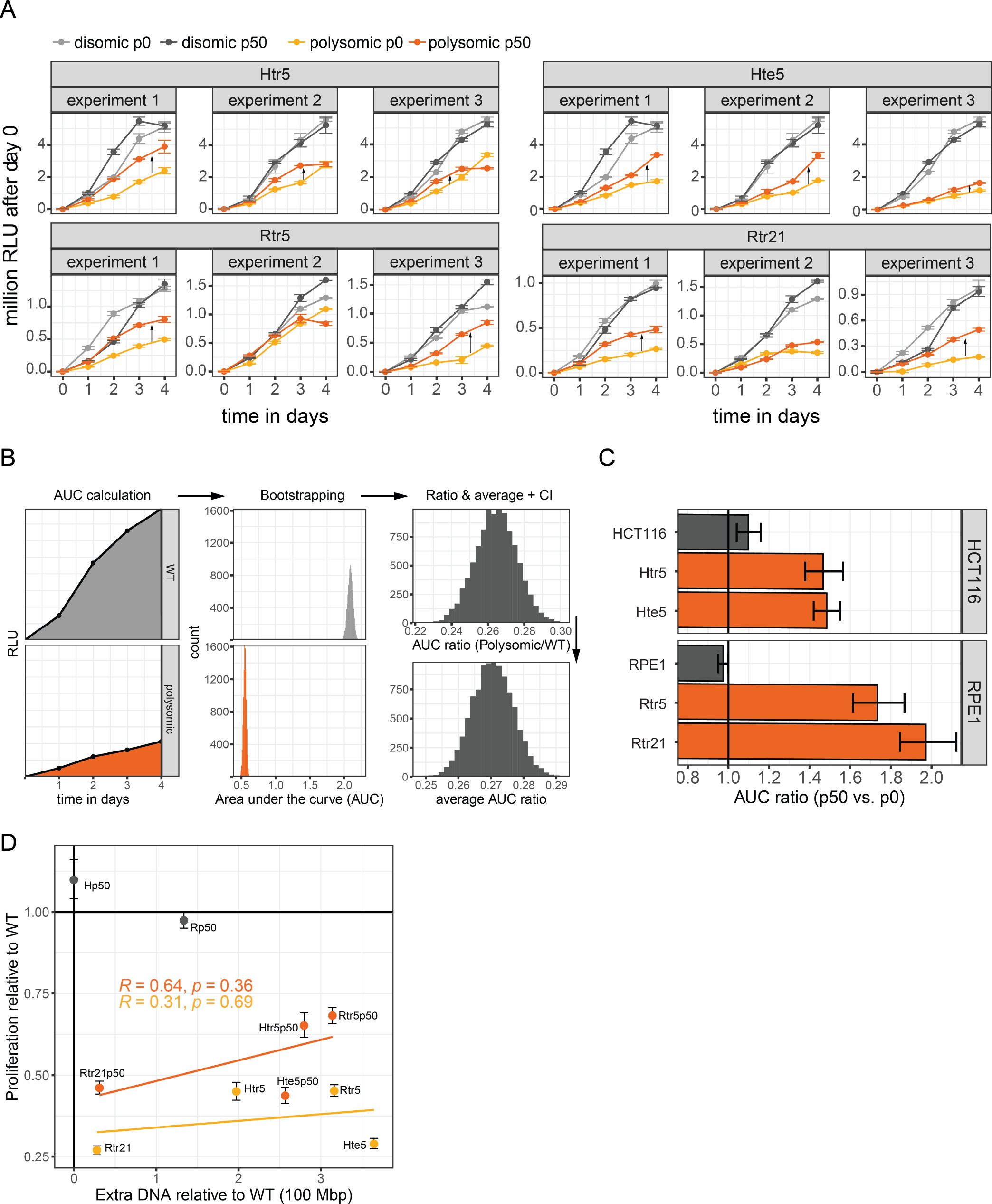
Evaluation of the proliferation changes in the polysomic cells after *in vitro* evolution. **A**. Growth curves from all experiments of population growth of the cell lines before and after *in vitro* evolution evaluated by MTT assay and normalized to the time point 0. Points represent mean relative light units (RLU) with SEM. **B**. Schematic of growth curve analysis pipeline consisting of calculating, bootstrapping, normalizing and averaging the areas under the curve (AUC) per experiment (N = 3). **C**. Ratios of AUCs between passage 50 and passage 0 cell lines. Bars represent empirical 95% confidence internals (10000 bootstrap samples). **D**. Correlation of the change in proliferation and change in total DNA relative to unevolved WT. Values depict Pearson correlation coefficient and p-value for p0 and p50 polysomic cell lines, bars represent empirical 95% confidence intervals.

**Figure S2.**
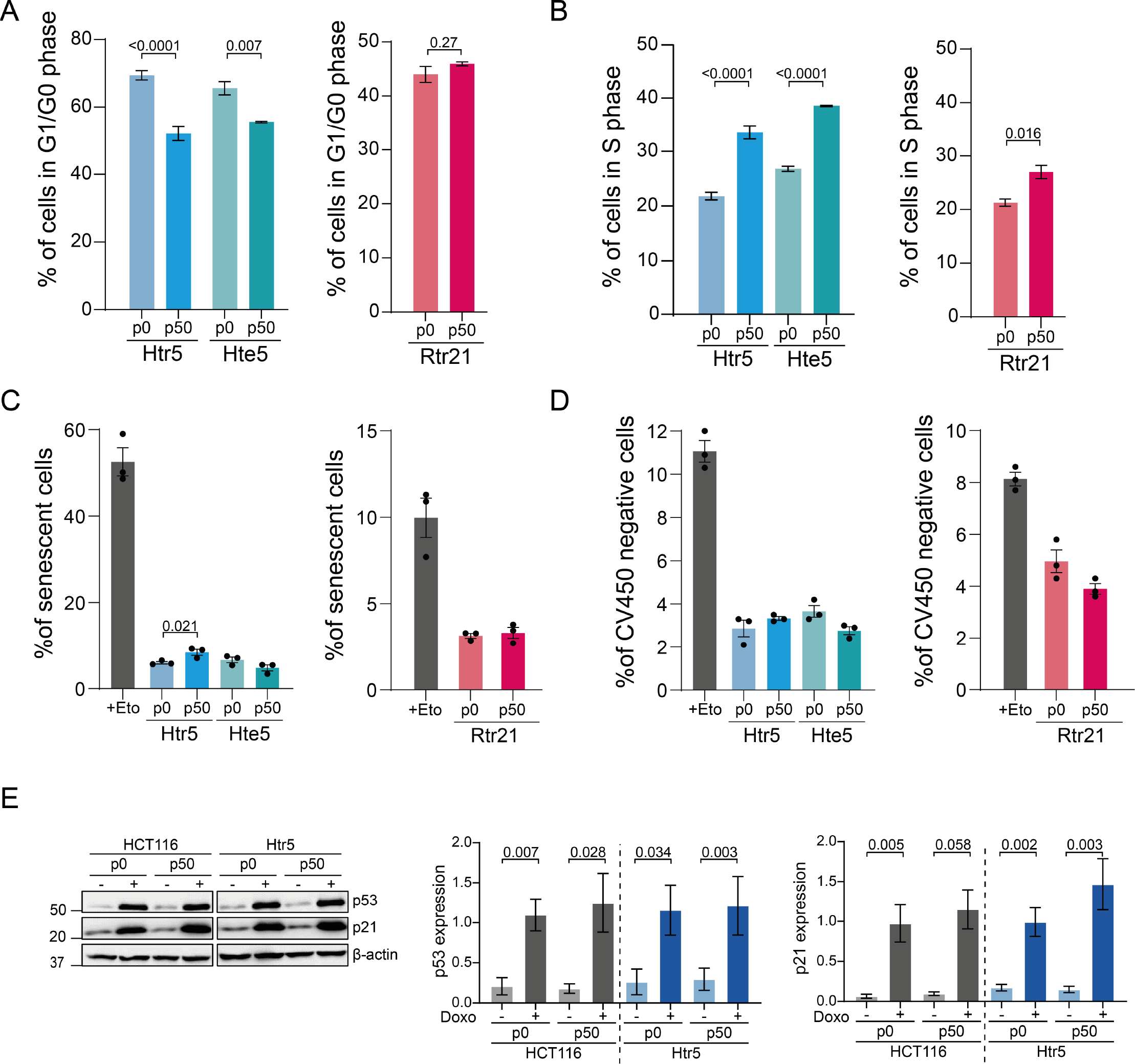
Evaluation of the cell cycle, cell death, senescence and p53 signaling in the polysomic cells after *in vitro* evolution. **A**. Relative proportion of cells in G1/G0 phase in polysomic cell populations before and after *in vitro* evolution determined with flow cytometry using EdU incorporation and DAPI staining (N: 3 - 9). **B**. Relative proportion of cells in S phase in polysomic cells before and after evolution determined with flow cytometry using EdU incorporation and DAPI staining (N: 3 - 9). **C**. Senescence in polysomic cells before and after *in vitro* evolution (N: 3). **D**. Proportion of CV450 viability dye negative cells in polysomic cell populations before and after *in vitro* evolution (N: 3). **E**. Representative immunoblot of p53 and p21 before and after *in vitro* evolution with and without doxorubicin treatment and quantifications. Doxorubicin treatment was performed with 1 μM overnight (N: 4 - 5). Data information: Mean with SEM is shown in all bar plots. In all plots unpaired Student’s t-tests were used for statistical evaluation.

**Figure S3.**
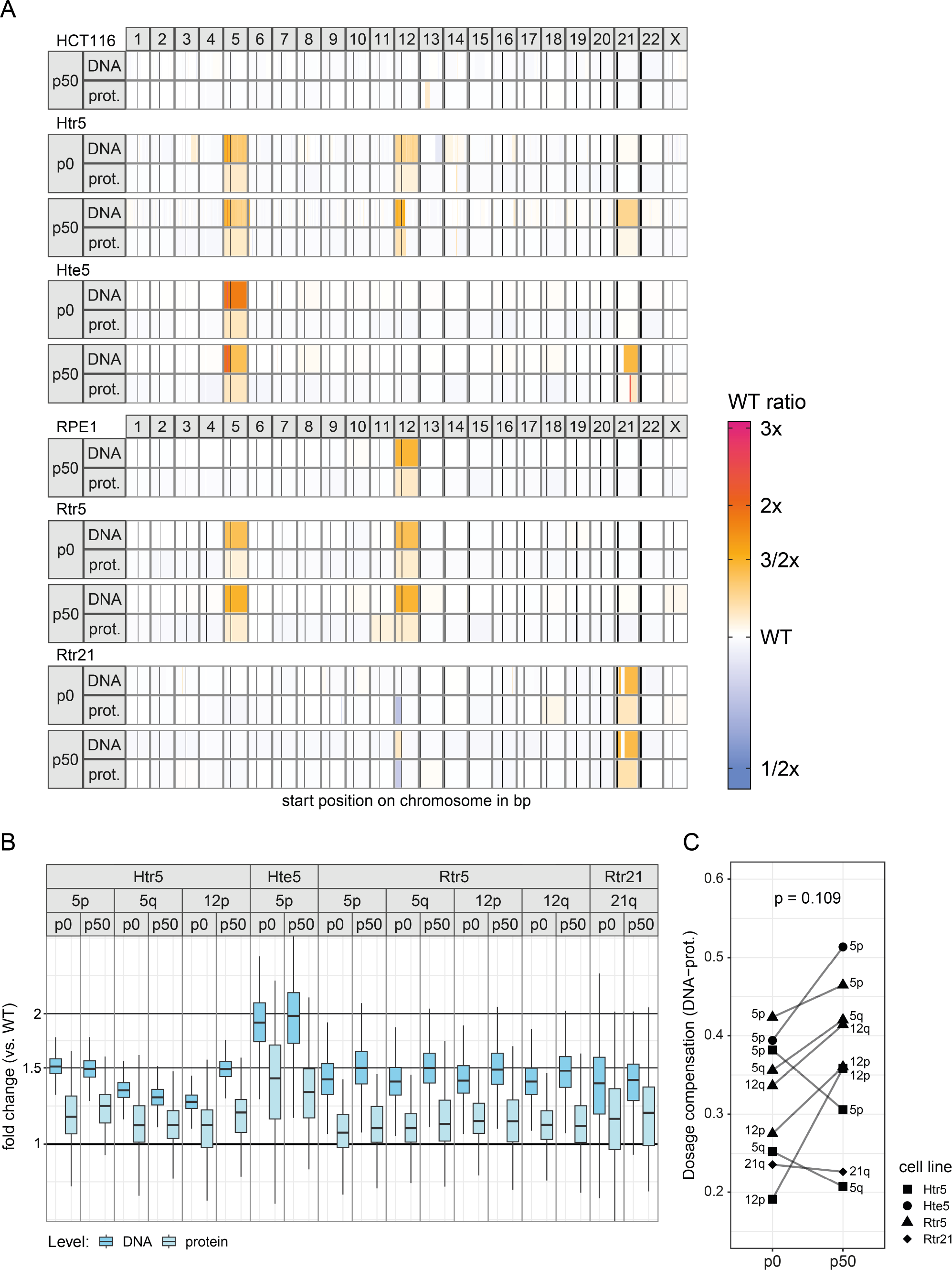
Genome and proteome changes before and after *in vitro* evolution. **A**. DNA copy number ratios and protein abundance fold changes relative to parental, unevolved WT for each cell line (row) and chromosome (column) as grouped by circular binary segmentation. Vertical bars within chromosomes depict the location of centromeres. **B**. Fold change in DNA copy numbers and protein abundances relative to parental, unevolved WT for supernumerary chromosome arms, which were not lost during *in vitro* evolution. **C**. Chromosome arm-level dosage compensation, calculated as the difference between average DNA copy number and protein abundance fold changes, before (p0) and after (p50) *in vitro* evolution.

**Figure S4.**
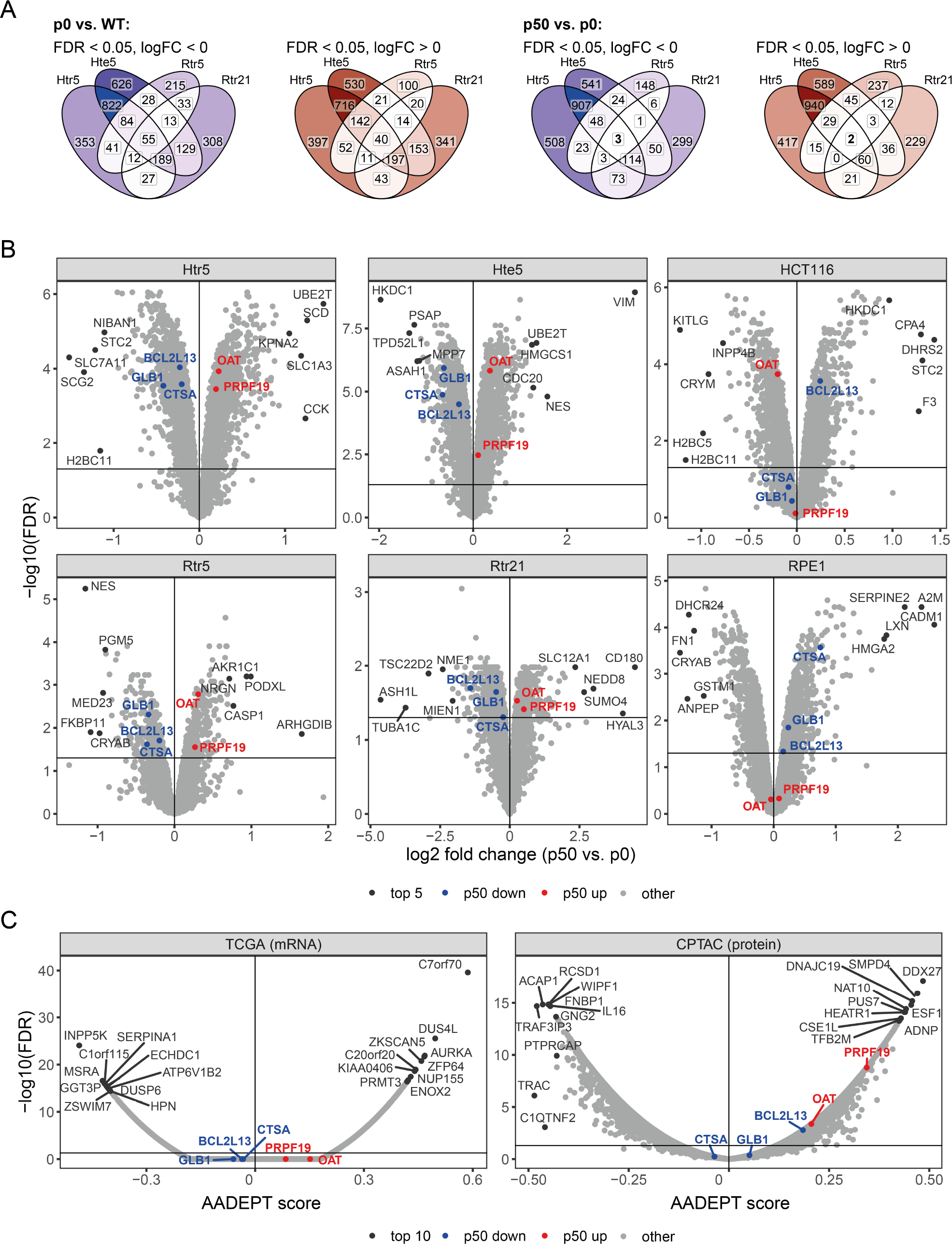
Gene expression changes of individual proteins after *in vitro* evolution and during tumorigenesis. **A**. Venn diagrams visualizing the overlap in significantly over- and underabundant proteins (FDR < 0.05) for cell line comparisons between polysomic cell lines and the parental WT (p0 vs. WT) and between polysomic cell lines before and after evolution (p50 vs. p0). **B**. Volcano plots showing changes in relative protein abundance after *in vitro* evolution of polysomic and WT cell lines. Proteins significantly over- (red) and underabundant (blue) in all adapted polysomic cell lines are highlighted next to the top 5 most over- and under-abundant proteins per individual comparison. **C**. Volcano plots of AADEPT scores for all measured genes on transcript-(TCGA) and protein-level (CPTAC). Genes with top 10 lowest and highest AADEPT scores as well as genes of proteins in the model cell line overlap are highlighted.

**Figure S5.**
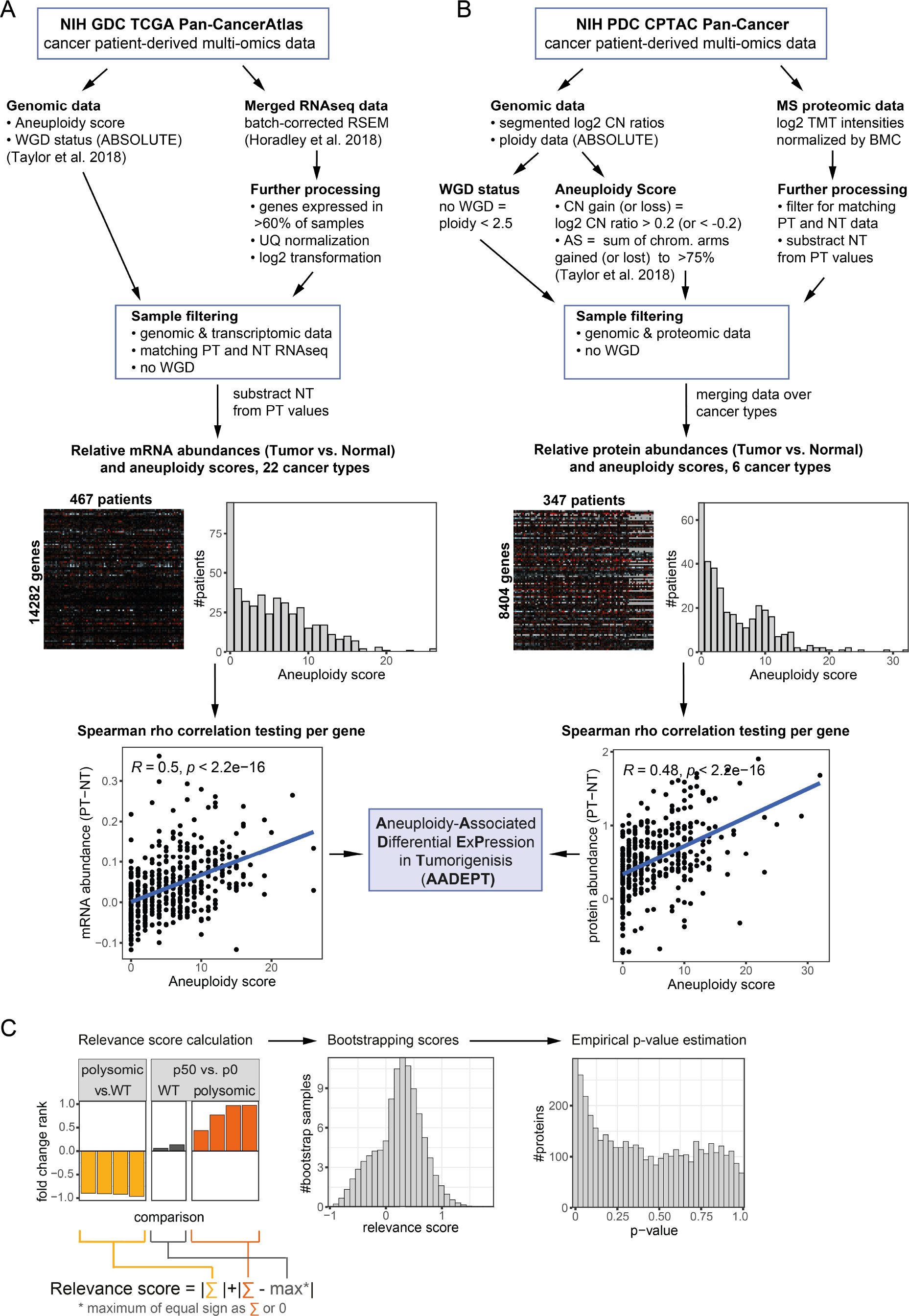
Schematic depiction of the strategy to calculate AADEPT scores and the aneuploidy relevance score. **A**. Derivation of AADEPT scores as Spearman correlation coefficients between the degree in aneuploidy and the change in gene expression in cancer patients (primary tumor vs. normal tissue) from multi-omics data of TCGA Pan-Cancer Atlas and **B**. from the Clinical Proteomic Tumor Analysis Consortium. **C**. Scoring of proteins based on relevant expression change patterns in our evolving polysomic model cell lines and deriving empirical p-values by bootstrapping the calculated scores. The relevance score is defined as the absolute sum of protein abundance fold change ranks (scaled from -1 to 1) of comparisons between evolved and unevolved polysomic cell lines as well as unevolved polysomic cell lines and their respective wild type. Congruent changes in wild type cells with *in vitro* evolution are penalized by subtracting the maximum fold change rank of equal sign. Data information: GDC = Genomic Data Commons, PDC = Proteomic Data Commons, TCGA = The Cancer Genome Atlas, CPTAC = Clinical Proteomic Tumor Analysis Consortium, BMC = Baylor College of Medicine, AS = aneuploidy score, WGD = whole-genome doubling, PT = primary tumor, NT = normal tissue, CN = copy number, UQ = upper-quartile.

**Figure S6.**
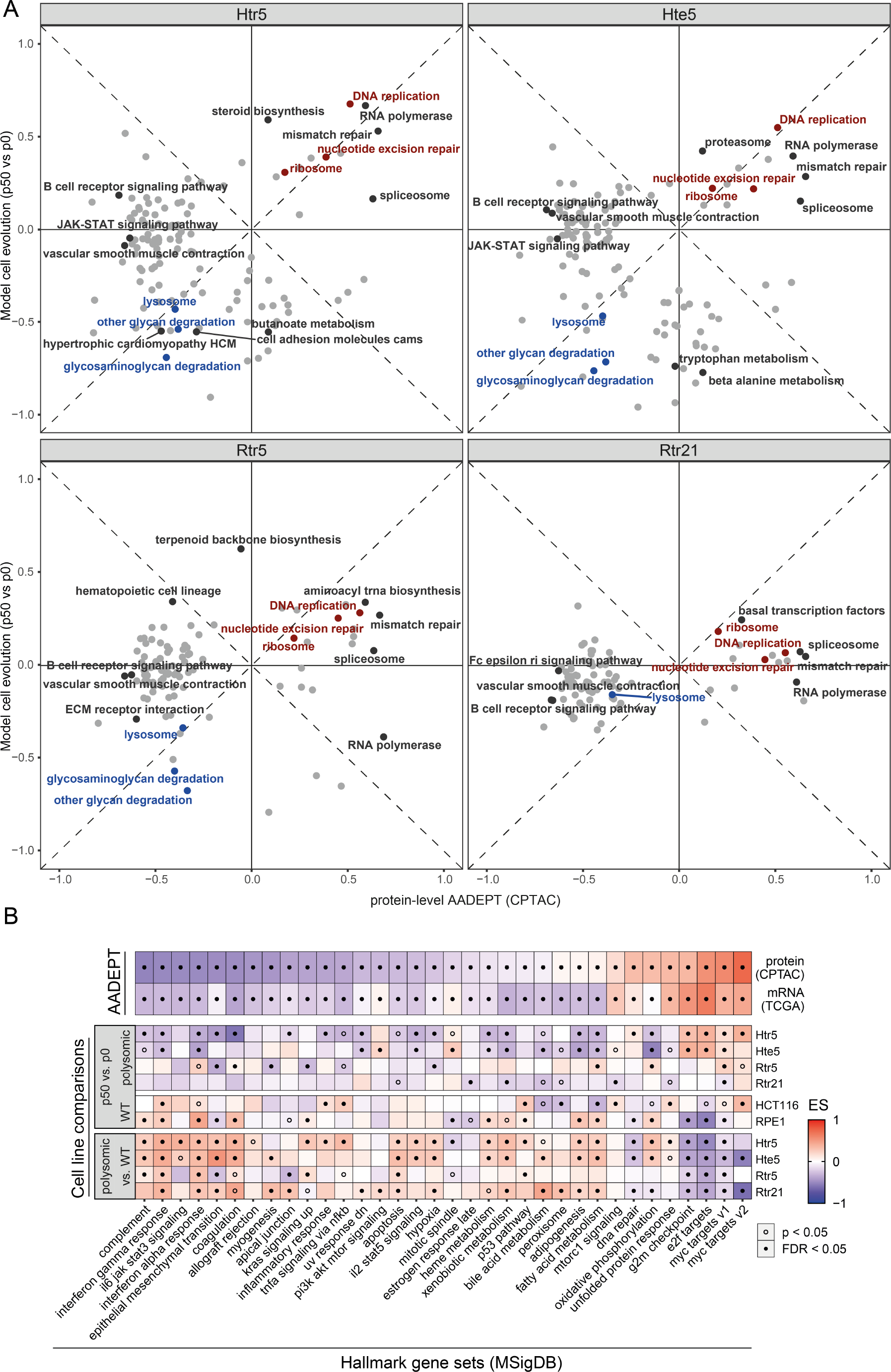
Gene expression changes of individual pathways after *in vitro* evolution and during tumorigenesis. **A**. 2D enrichment of KEGG pathways comparing protein-level AADEPT scores (x-axis) with fold changes in all polysomic cell lines after adaptation (y-axis), showing enrichment scores for pathways with FDR < 0.05 and highlighting those with shared enrichment in adapted polysomic cell lines (blue - negative, red - positive) as well as those with top 3 enrichment scores in both direction of either axis and with more than 10 measured proteins. **B**. Enrichment scores of Hallmark gene sets for AADEPT scores (above) and for protein abundance changes between cell line comparisons (below). Hallmark gene sets with FDR < 0.05 in a 2D enrichment analysis of transcript- and protein-level AADEPT scores are shown and sorted by degree of enrichment with protein-level AADEPT scores. The statistical significance of enrichment in individual comparisons is indicated by the points within each tile, p-value < 0.05, FDR < 0.05.

**Figure S7.**
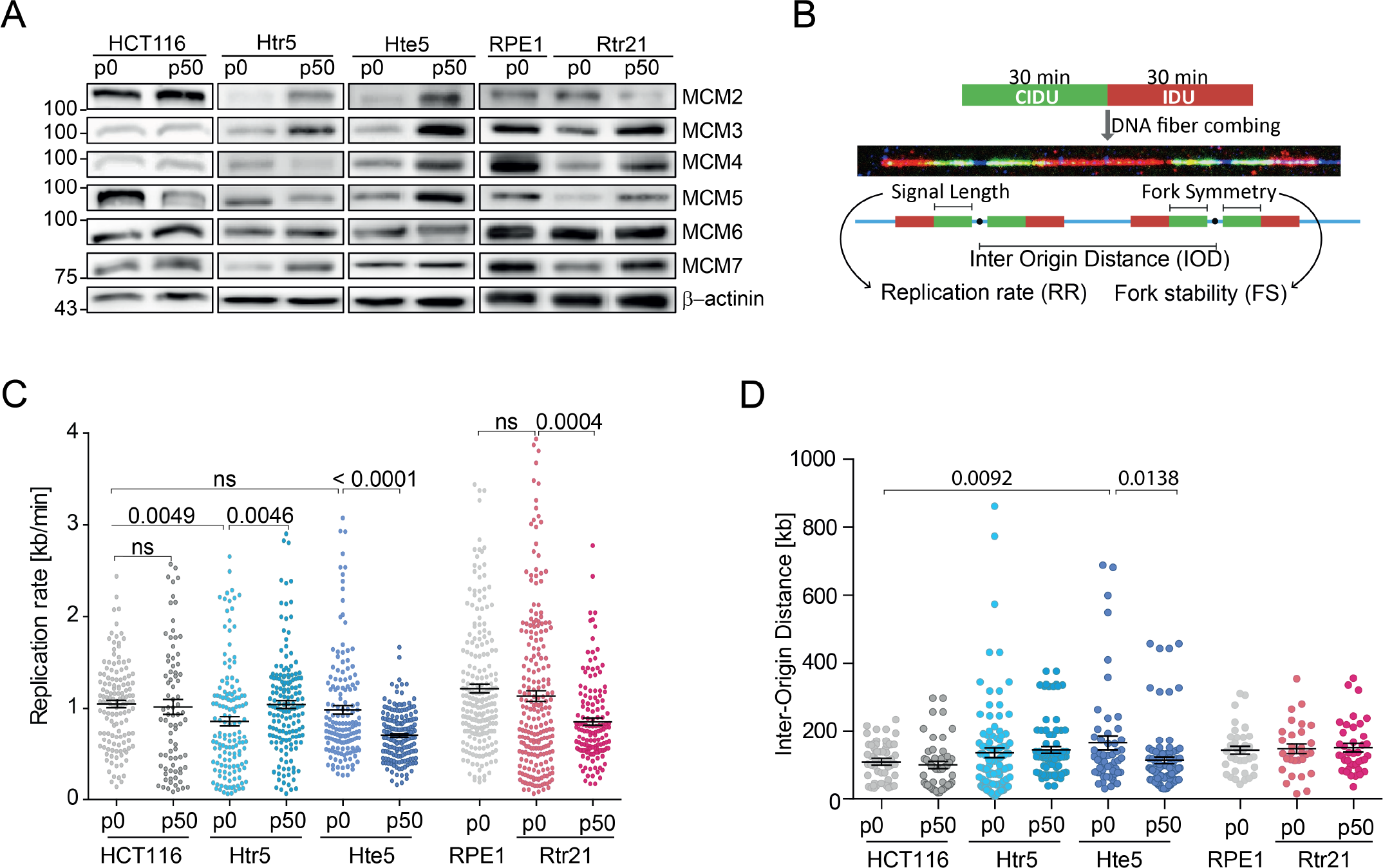
Changes in replication after *in vitro* evolution. **A**. Extended version of representative immunoblot of MCM2-7 in model cell lines before and after *in vitro* evolution. **B**. Schematic depiction of the replication dynamics analysis. **C**. Replication rate of the model cell lines before and after evolution. **D**. Inter-origin distance in model aneuploid cells before and after evolution. Data information: Mean with SEM is shown in all experiments. Unpaired Student’s t-tests were used for statistical evaluation.

**Figure S8.**
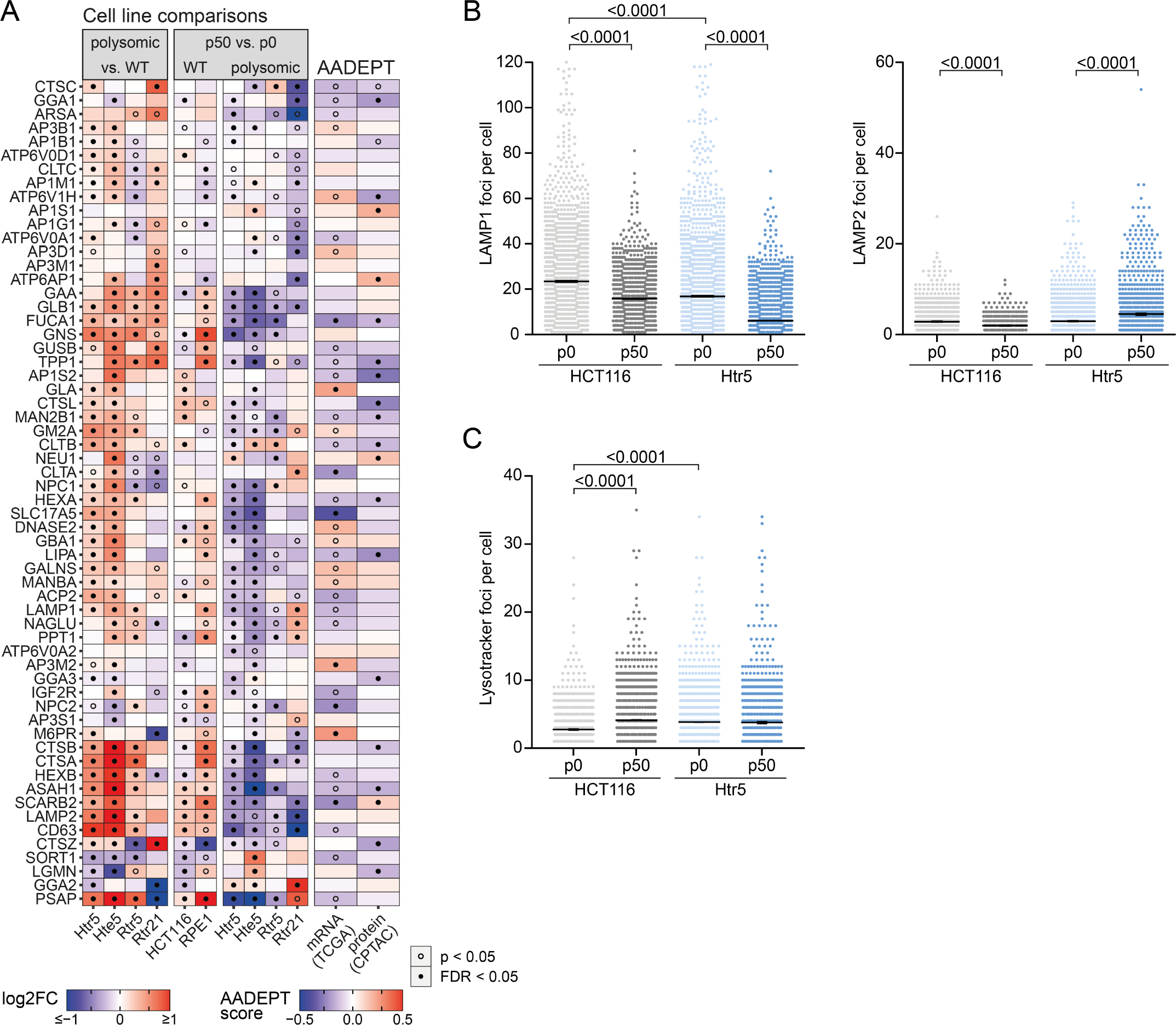
Changes in lysosome and lysosome degradation pathways and phenotypes after *in vitro* evolution. **A**. Expression changes of lysosomal proteins (KEGG) after chromosome gain (polysomic vs. WT) and after *in vitro* evolution (p50 vs. p0) and their corresponding AADEPT scores (as in Fig. 2E). **B**. Quantification of immunofluorescence intensity of LAMP1 (left) and LAMP2 (right). Scatter plot of foci per cell (N ≥ 3600). For the LAMP1 analysis values greater than 120 foci per cell were excluded since they were most likely artifacts of high background. Mean with SEM. (N ≥ 3600, three independent experiments with three technical replicates each). **C**. Quantification of cells with lysotracker foci. Scatter plot of foci per cell. Mean with SEM. (N ≥ 2400, three independent experiments with three technical replicates each). Significance was analyzed using Student’s t-test.

**Figure S9.**
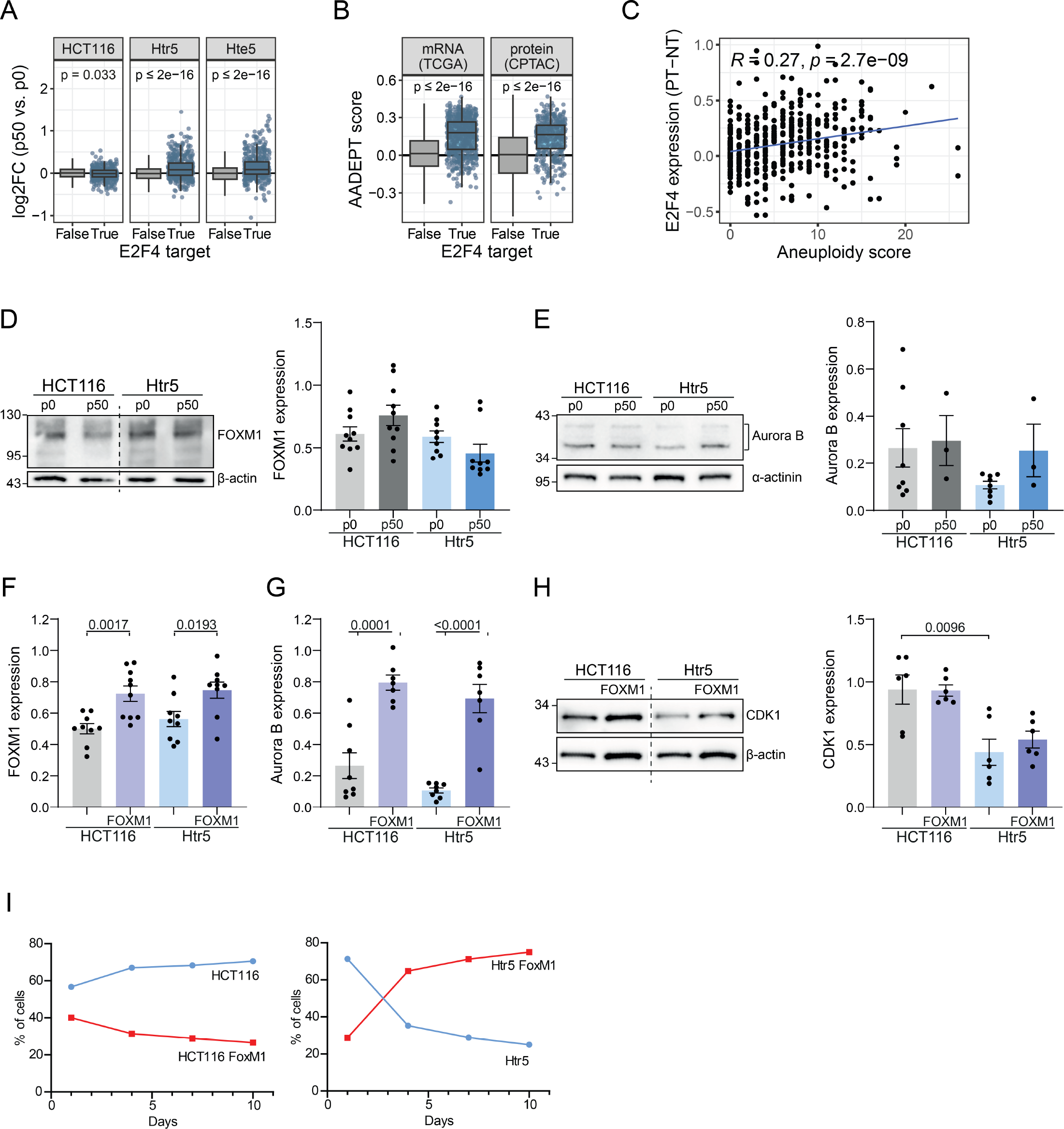
FOXM1 and E2F4 dependent changes after *in vitro* evolution and in cancer. **A**. Protein abundance fold changes of all E2F4 targets in evolved model cell lines, tested against all other proteins using Welch’s t-tests (N: 484). **B**. Transcript-(TCGA) and protein-level (CPTAC) AADEPT scores of E2F4 target genes tested against all other proteins using Welch’s t-tests (N: 648, N: 406). **C**. Spearman correlation coefficient between TCGA patient (N = 467) tumor aneuploidy score and E2F4 gene expression relative to normal tissue. **D**. Representative immunoblot of FOXM1 expression and quantification before and after evolution (N: 9 - 10). **E**. Representative immunoblot of Aurora B kinase expression and quantification before and after evolution (N: 3 - 8). **F**. Quantification of immunoblotting of FOXM1 overexpression (N: 9 - 10). **G**. Quantification of immunoblotting of Aurora kinase B in FOXM1 overexpressing cells (N: 7 - 8). **H**. Representative immunoblot of CDK1 in FOXM1 overexpressing cells and quantification (N: 6). **I**. Quantification of the RFP and BFP positive cell fraction in competition assay. Representative experiment. Data information: Mean with SEM is shown for all bar plots. In all plots unpaired Student’s t-tests were used for statistical evaluation.

**Figure S10.**
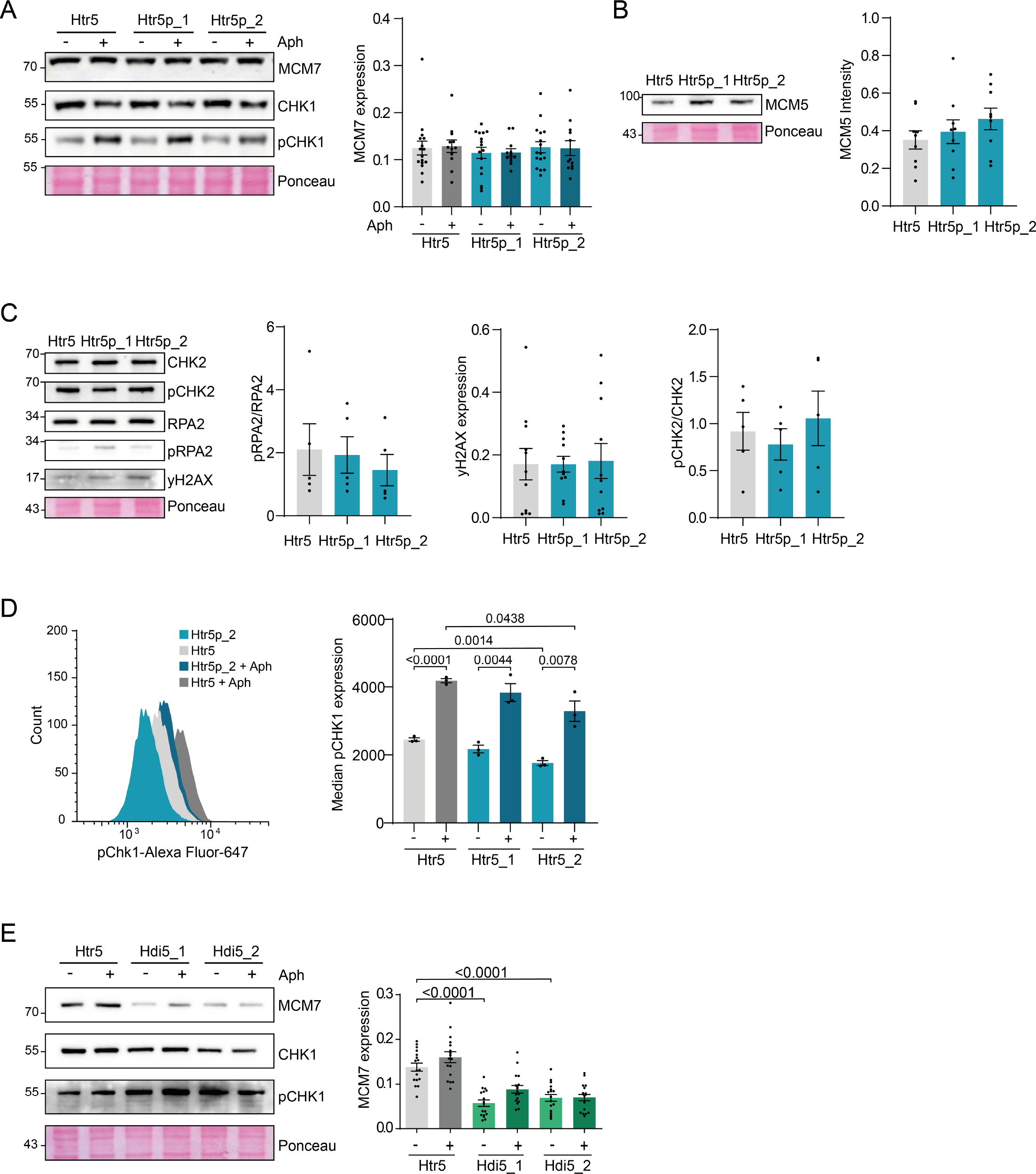
Effect of chromosome 5 arms on replication stress-related protein expression. **A**. Representative immunoblot of MCM7 and pCHK1(S345)/CHK1 expression in Htr5 and Htr5p cell lines with MCM7 quantification. **B**. Representative immunoblot of MCM5 expression with quantification. **C**. Representative immunoblot of CHK2, pCHK2, RPA2, pRPA2 (S33), and yH2AX with respective quantifications. **D**. Representative histograms of pCHK1 flow cytometry analysis with quantification. **E**. Representative immunoblot of MCM7, CHK1, and pCHK1(S345) expression in Htr5 and Hdi5 cell lines with MCM7 quantification. All experiments were performed in at least three biological replicates with 1-6 technical replicates each. Data information: Mean with SEM is shown for all bar plots. In all plots unpaired Student’s t-tests were used for statistical evaluation.

